# A Connectivity-based Psychometric Prediction Framework for Brain-behavior Relationship Studies

**DOI:** 10.1101/2020.01.15.907642

**Authors:** Jianxiao Wu, Simon B. Eickhoff, Felix Hoffstaedter, Kaustubh R. Patil, Holger Schwender, B.T. Thomas Yeo, Sarah Genon

**Affiliations:** Institute for Systems Neuroscience, Medical Faculty, Heinrich-Heine University Düsseldorf, Germany; Institute of Neuroscience and Medicine, Brain & Behaviour (INM-7), Research Center Jülich, Jülich, Germany; Mathematical Institute, Heinrich-Heine University Düsseldorf, Düsseldorf, Germany; Department of Electrical and Computer Engineering, ASTAR-NUS Clinical Imaging Research Centre, Singapore Institute for Neurotechnology and Memory Networks Program, National University of Singapore, Singapore City, Singapore; Athinoula A. Martinos Center for Biomedical Imaging, Massachusetts General Hospital, Charlestown, Massachusetts; Center for Cognitive Neuroscience, Duke-NUS Medical School, Singapore City, Singapore

**Keywords:** brain-behavior relationships, behavior prediction, human connectome project, machine learning, resting state functional connectivity

## Abstract

The recent availability of population-based studies with neuroimaging and behavioral measurements opens promising perspectives to investigate the relationships between interindividual variability in brain regions’ connectivity and behavioral phenotypes. However, the multivariate nature of connectivity-based prediction model severely limits the insight into brain-behavior patterns for neuroscience. To address this issue, we propose a connectivity-based psychometric prediction framework based on individual regions’ connectivity profiles. We first illustrate two main applications: 1) single brain region’s predictive power for a range of psychometric variables, and 2) single psychometric variable’s predictive power variation across brain region. We compare the patterns of brain-behavior provided by these approaches to the brain-behavior relationships from activation approaches. Then, capitalizing on the increased transparency of our approach, we demonstrate how the influence of various data processing and analyses can directly influence the patterns of brain-behavior relationships, as well as the unique insight into brain-behavior relationships offered by this approach.

## 1. Introduction

The relationships between brain regions and behavioral functions is a fundamental question for neuroscience. These relationships can be investigated by relating interindividual variability in brain regions’ connectivity to interindividual differences in behavioral performance (that is, to psychometric data). The recent availability of population-based neuroimaging datasets with extensive psychometric characterization (Van Essen et al. 2013; Caspers et al. 2014) opens promising perspectives to investigate the relationships between brain regions’ connectivity and behavior. In particular, many studies have shown that individual profile of functional connectivity (FC) between brain regions can predict individual scores on psychometric variables, including cognitive measures such as fluid intelligence, as well as personality traits, such as openness (Finn et al. 2015; Rosenberg et al. 2016; Smith et al. 2016; Noble et al. 2017; Dubois et al. 2018; Li et al. 2019; Pervaiz et al. 2020). Given the potential of these approaches for cognitive and clinical neuroscience, developing a scientifically valid and useful connectivity-based framework for investigating brain-behavior relationships is a crucial objective for the neuroimaging community.

In recent studies, important attention has been given to improving the predictive model performance, which is evaluated by comparing the predicted behavioral score with the observed one. For instance, the predicted and actual fluid intelligence values show, across many different studies, a Pearson correlation coefficient r of around 0.25 (Smith et al. 2016; Dubois et al. 2018; Li et al. 2019; Pervaiz et al. 2020), while a correlation of around r = 0.11 could be achieved on average across 58 different measures covering different domains (Li et al. 2019). Such relatively low model performance suggest that the field is not yet mature and that predicting interindividual variability in behavioral performance from interindividual variability in FC remains particularly challenging. Furthermore, in a cognitive and clinical neuroscience framework, not only the prediction performance matters, but also the neurobiological validity of the model and relatedly, its interpretability, raising one main issue of the current state of the art. Frequently, researchers try to interpret the model with some type of post-hoc evaluation, looking at the brain connectivity features (i.e. region-to-region connectivity values) seemingly playing important roles in the prediction. The relative relevance of the features is often derived from the weights assigned by the regression algorithms. Nevertheless, since prediction models often enlist backward (or discriminative) models, such interpretations can be drastically misleading as the relative magnitude of these weights does not reflect the magnitude of the regions’ association with the given psychometric variable, and large weights may be assigned to features which are actually unrelated to any brain process of interest (Haufe et al. 2014). The difficulty of interpretations could also be illustrated with the example, shown in figure 1, of average weights assigned during prediction of fluid intelligence by support vector regression (SVR; Boser et al. 1992; Cortes and Vapnik 1995) and elastic nets (EN; Zou and Hastie 2005) respectively. The small number of highlighted connectivity edges suggests that interpretations based on only these edges could be unrepresentative. The inconsistency of weight assignment across regression algorithms also suggests the lack of reliability for any interpretations. To address this issue, we here propose a connectivity-based psychometric prediction (CBPP) framework based on individual region’s connectivity profile. Such an approach is performed by evaluating a machine learning model predicting psychometric data from FC independently for each brain region (see Chen et al. 2020 for a previous application in classification). The prediction performance of psychometric data of each independent brain region’s model can then be used as an estimator of the relationship between the region’s connectivity profile and the measured behavioral function.

**Figure 1.**
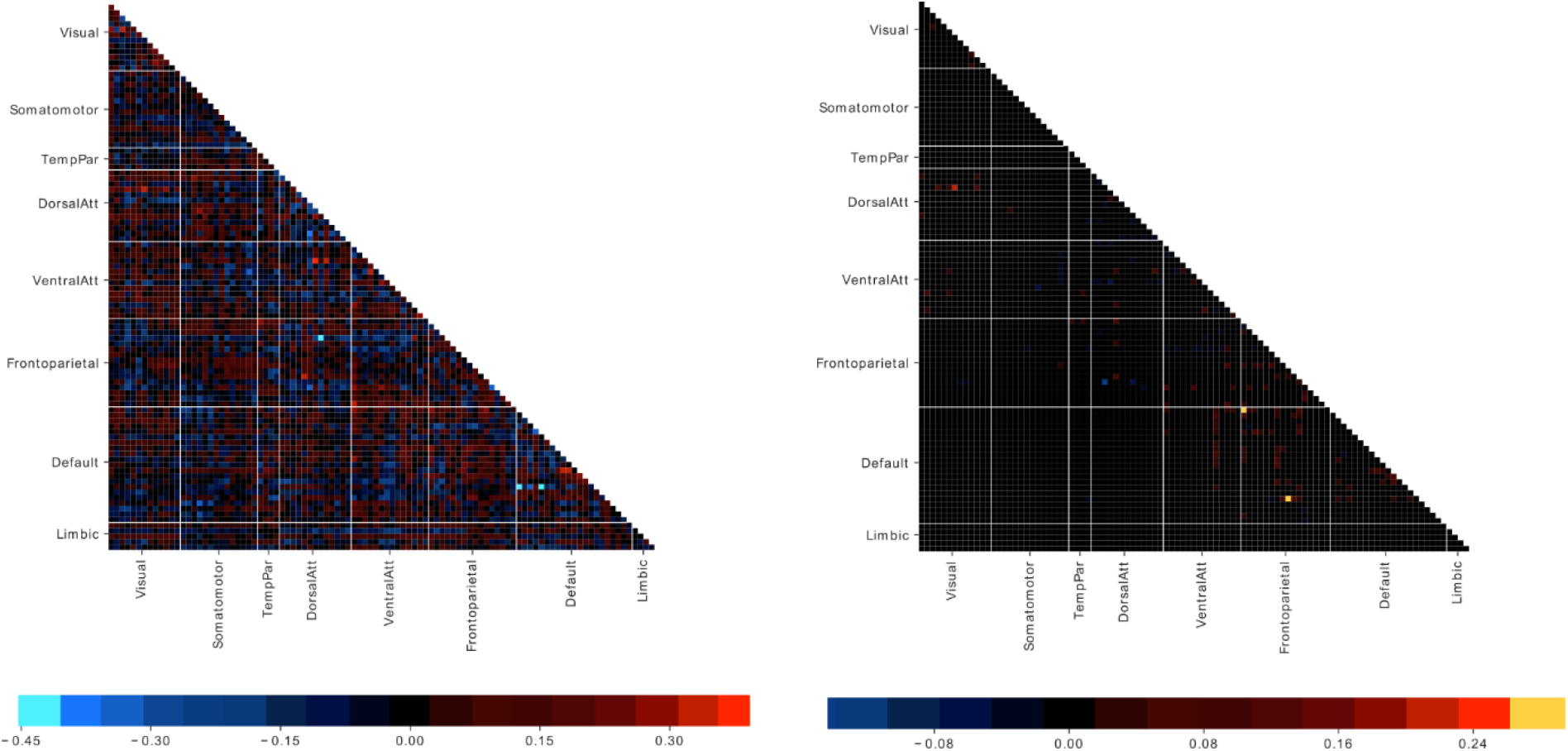
Weights of connectivity features assigned by SVR (left) and EN (right) for prediction of fluid intelligence, computed on the FIX-processed data from the Human Connectome Project described in the ‘Data and preprocessing’ section. Color represents the average weight value across one run of 10-fold cross-validation.

The relationship between brain functional connectivity and behavior is multivariate with a level of complexity that prevents any one-to-one mapping between brain region and behavioral function (Genon et al. 2018). Many “global” (whole-brain connectivity or network-based connectivity) approaches aim to account for this complexity in the prediction of behavior (Kong et al. 2019; Kashyap et al. 2019; Pervaiz et al. 2020). A local approach, while obviously simplifying the complexity of brain function (Horien et al. 2019), offers additional insights into individual brain regions of interest. Such insights are necessary for a progressive unraveling of the brain-behavior relationship and hence the investigations of the neurobiological validity of the prediction. For instance, if parcels in visual areas are the best at predicting abstract reasoning performance, one could expect that the model is partly driven by some confounding factors such as the posterior brain shape. Furthermore, from such insights, a systematic examination of the influence of different factors on the prediction model can be conducted. One can hence for instance, investigate how controlling for brain shape estimators influences the pattern of relationships between brain regions and behavioral variables. Finally, from a more clinical standpoint, a region-based approach can offer specific insights into patterns of brain regions’ dysfunction and behavioral symptoms. In sum, a region-based approach brings a complementary insight to currently used whole-brain and network-based approaches by the transparency it offers and hence provides a new insight into brain-behavior relationships.

In the current study we first examined the relationships between brain region connectivity and psychometric variables by looking at the prediction profile across psychometric variables of a given region, followed by depicting, for a given psychometric score, the profile of distribution of prediction performance across brain regions. Then, in order to better understand how methodological choices can affect our study of brain-behavior relationships, we illustrated the influence of confounds that are not systematically taken into account in the literature, such as brain size. We also examined whether sophisticated denoising of resting-state functional connectivity (RSFC) provides a clarified picture of brain-behavior patterns or rather appears to remove relevant signal. Lastly, we compared our approach with a post-hoc evaluation of the nodes or edges playing a role in the predictions (as shown in figure 1). Following this first demonstration of the insight that our approach can bring, we will discuss our results and the related open challenges.

## 2. Materials and Methods

### 2.1. Data and preprocessing

In this study, for extensive evaluation on high quality data, the 1200 Subjects Data Release of the Human Connectome Project (HCP) data was used (Van Essen et al. 2013). Subjects were young healthy adults (aged 22-37), from families with twins and non-twin siblings. Imaging data were acquired using a customized Siemens 3T Skyra. Each subject visited in two consecutive days, during each of which two rs-fMRI runs were acquired using different phase encodings, left-right and right-left. Each run is 1200 frames (14.4 min) in length, with a repetition time (TR) of 720 ms. We only considered subjects with all 4 runs completed.

All the raw rs-fMRI data from HCP were preprocessed by the HCP Minimal Processing Pipelines (Glasser et al. 2013), which include motion correction, gradient nonlinearity distortion correction, EPI distortion correction, coregistration to T1-weighted images and normalization to the MNI152 space. We refer to the resulting data as “minimally processed”. Further cleaning of noise in HCP data was done using ICA-FIX. Components were first identified using independent component analysis (ICA); a classifier (FIX) trained on HCP data was then applied to remove artifactual components (Smith et al. 2013; Salimi-Khorshidi et al. 2014; Griffanti et al. 2014). We refer to these data as “FIX”. To further investigate the impact of global signal regression (GSR), we regressed the cortical global signal, which is the average signal across all cortical vertices, and its temporal derivative (Power et al. 2018; Li et al. 2019) from the FIX data. These data are then referred to as “FIX+GSR”. We obtained surface data in the fsLR space for these three types of preprocessing strategies (N=928 for ‘minimally processed’, N=923 for ‘FIX’ and ‘FIX+GSR’data). For the volumetric data, we only used the existing FIX denoised data (N = 931). Additionally, we applied nuisance regression to control for the 24 motion parameters, white matter (WM) and cerebrospinal fluid (CSF) signals, and their derivatives. We refer to these data as “FIX+WM/CSF”.

For most subjects of the HCP cohort, a large number of psychological measures were collected through tests and questionnaires across a broad range of psychological domains including sensory, motor, cognition, emotion, affect and personality. Additionally, task performance scores were also available from task-fMRI sessions. We selected 40 representative psychometric variables after inspecting all variables distribution for roughly normal distribution. See Table S1 for the complete list of variables selection.

Psychometric data are typically related to demographical factors such as age and gender. In addition, the relationships between psychometric and neuroimaging data can be partially mediated by factors such as head size, handedness and seasonality. To ensure strict control on the confounding effects of these latter variables, we not only regressed out the first level effect of brain size, sex and age but also included the secondary term of age and the interaction between age and sex, in line with the HCP MegaTrawl analysis (Smith et al. 2016). Accordingly, our ‘standard’ confound controlling method involved regressing out 9 confounding variables from the psychometric variables: sex (*Gender*), age (*Age_in_Yrs*), age^2^, sex*age, sex*age^2^, handedness (*Handedness*), brain size (*FS_BrainSeg_Vol*), intracranial volume (ICV; *FS_IntraCranial_Vol*) and acquisition quarter (*Acquisition*).

Differing from some previous work, we did not include motion-related confounding variables, such as framewise displacement (FD) and DVARS. First, the effects of these confounds on the resting-state fMRI data should be largely reduced following motion correction and FIX denoising. Second, the correlation between these confounds and the psychometric variables are rather low, especially after controlling for the above 9 confounding variables (see Table S2 and S3). Finally, from a conceptual standpoint, while it can be assumed that motion-related effects could influence association between whole-brain patterns and psychometric data, there is no reason to expect head motion effect in specific brain regions that would hence specifically affect the predictive pattern of these regions. Accordingly, we did not observe any major effect of these confounds on parcel-wise psychometric profiles (see a comparison in figure S2). As a result, FD and DVARS were not included as confounding variables, since they would not affect our conclusions.

### 2.2. Preliminary evaluation

Preliminary to the development of our region-wise prediction approach, we performed an extensive preliminary assessment of the general CBPP framework based on a global approach, that is, based on whole-brain connectivity information. The general workflow of the CBPP framework follows standard protocols (Shen et al. 2017) and is illustrated in Figure S1. At each step of the framework, multiple approaches and various parameter settings could be considered. In our evaluation, we mainly considered previously used approaches, summarized in Table S2. Figure 2 shows the different approaches considered at each step in our implementation of whole-brain CBPP. In total, 96 combinations of approaches were evaluated with ‘standard’ confound controlling approach. We used the surface data from HCP, including all 3 different preprocessing strategies (‘minimally processed’, ‘FIX’ and ‘FIX+GSR’).

**Figure 2.**
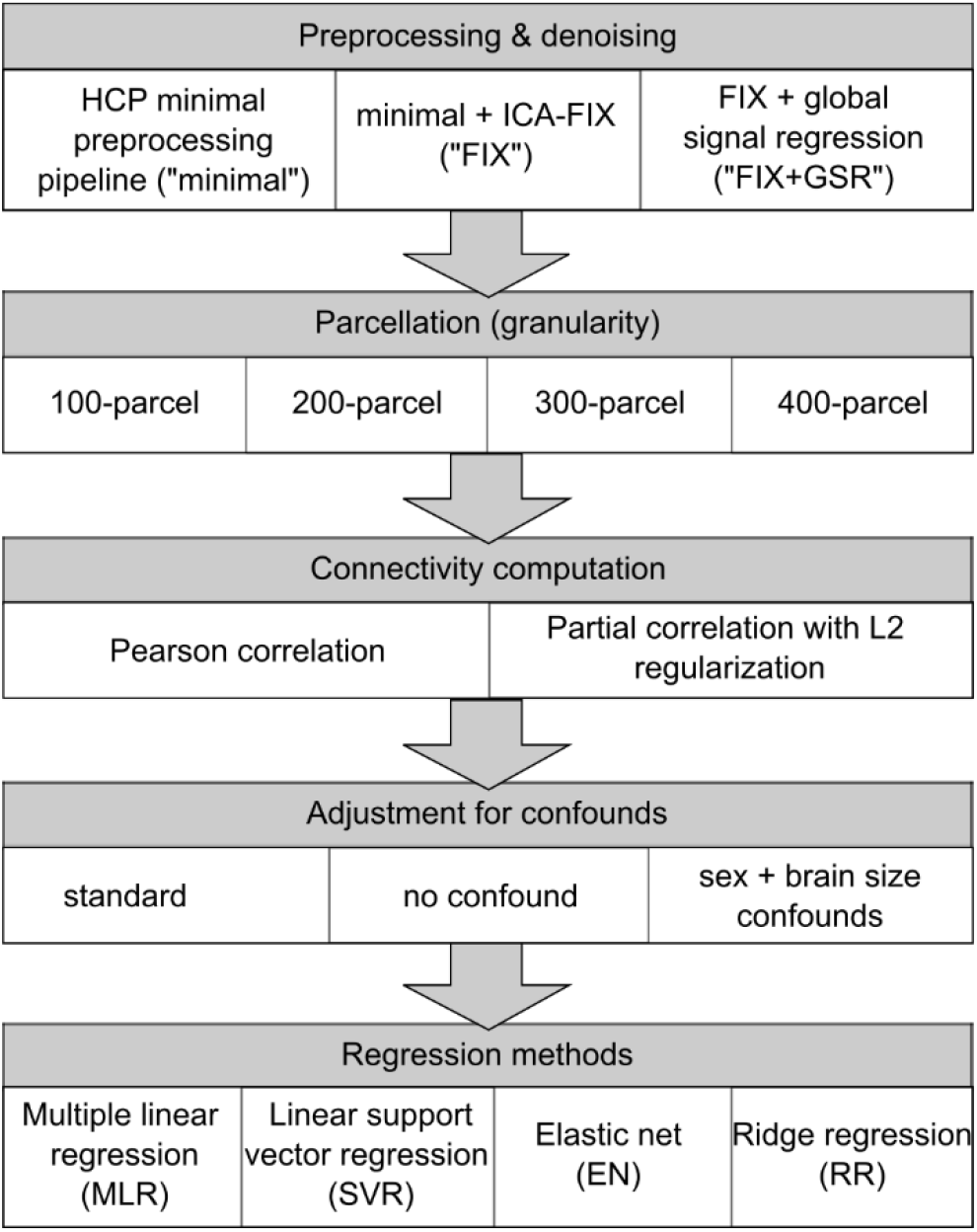
Approaches considered for each step in the CBPP framework.

CBPP studies usually capitalizes on whole-brain connectivity using atlases (brain parcellation) to summarize voxels data, by computing a mean time series for each parcel by averaging time series of all the vertices (or voxels) within that parcel. In our implementation, the Schaefer atlas (Schaefer et al. 2018) was utilized at 4 different granularity levels, with 100 parcels, 200 parcels, 300 parcels and 400 parcels respectively. The most commonly used functional atlases contain 200 to 400 parcels (Shen et al. 2013; Joliot et al. 2015; Gordon et al. 2016; Glasser et al. 2016). Nonetheless, in the recent HCP MegaTrawl, the highest psychometric prediction accuracy was achieved at lower dimensionality, with dimension d = 50, when partial correlation (with L2 regularization) was used for computing connectivity (Smith et al. 2016). Therefore, we used all available granularity up to 400-parcel level.

Functional connectivity is typically computed based on Pearson correlation; however, partial correlation is considered by several authors as more likely to reflect direct connections (Bijsterbosch et al. 2017; Lim et al. 2019). When using atlases, functional connectivity strengths are computed as the correlation coefficient between each pair of parcel’s average time series resulting in, for example, a 400×400 FC matrix at 400-parcel granularity. In our implementation, the Pearson correlation matrices were computed using the Matlab’s corrcoef function. The partial correlation matrices were computed using FSLNets’ nets_netmats function (http://fsl.fmrib.ox.ac.uk/fsl/fslwiki/FSLNets), with the default regularization coefficient for L2-norm ridge regression. The L2 regularization adds a quadratic term of the regression weights to the cost function, hence adding a constraint to the optimization. Such constraints help to prevent overfitting by shifting the best-fit solution on training set by an amount independent of the data. Since 4 runs were acquired for each HCP subject, the average connectivity matrix for each subject across runs was used as input features for the supervised learning model.

Commonly in literature, feature selection can be done before the prediction step to reduce computation time (Smith et al. 2016; Shen et al. 2017; Dubois et al. 2018; Kashyap et al. 2019). One popular way to do so is to examine Pearson correlation coefficients between elements of the connectivity matrix (i.e. edges) and the psychometric variables to predict and to select either connectivity elements with significant correlation (p < 0.01 or p < 0.05) or the top 50% of all the connectivity elements (i.e. 2475, 9950, 22425 and 39900 elements at 100-parcel, 200-parce, 300-parcel, and 400-parcel granularity respectively) as features for the subsequent prediction step. In our implementation, we selected the top 50% of all connectivity elements as features for the more computationally expensive algorithm, EN. As multiple linear regression (MLR) relies on least square solutions and was mostly used in previous studies for univariate prediction, it could not be expected to deal with higher feature dimensionality. Therefore, we only selected the top 500 of all connectivity elements as features for MLR in order to prevent overfitting. No feature selection was performed for linear SVR or ridge regression (RR), since the computation was not particularly expensive and, importantly, no overfitting was observed.

Finally, we focused on a selection of linear regression techniques popular in the neuroscience field: multiple linear regression (Matlab’s regress function; Chatterjee and Hadi 1986), linear SVR, EN, and RR. For each combination of approaches, we performed 10 repeats of 10-fold cross-validation. In each repeat, the subjects were divided into 10 folds. For each test fold, the regression model was estimated using the remaining 9 training folds. The corresponding prediction accuracies were then computed by applying the model to the test fold. The final prediction accuracy was measured as 1) the average Pearson correlation 2) inverse of the average normalized root mean squared deviation (nRMSD; Pineiro et al. 2008; Dubois et al. 2018) between the predicted and observed psychometric values, across all test set folds and all repeats. In particular, the inversed nRMSD values were calculated as:

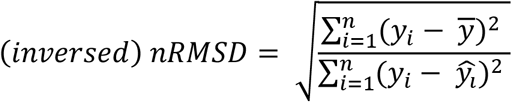

For all combinations of approaches tested in the preliminary evaluation, the difference in performance between combinations were tested with the corrected resampled t-test (Nadeau and Bengio 2003; Bouckaert and Frank 2004), which accounts for the dependency between test folds across repeats. The results were corrected for multiple comparisons using false discovery rate (FDR; Hochberg & Benjamini 1995) of q < 0.05.

We note that the comparisons between the different regression algorithms may not be generalizable to different frameworks and fully objective, since different feature selection methods were utilized for the different algorithms. These comparisons therefore do not provide general rankings or guidelines for the community. They here serve as a preliminary optimization of the methods on which our region-based framework builds.

### 2.3. Psychometric prediction

For the prediction step, linear SVR and RR were implemented using Matlab’s fitrlinear function, while EN was implemented using the glmnet package for Matlab (Qian et al. 2013). We note that LASSO (least absolute shrinkage and selection operator) is also widely used in neuroscience studies. However, since EN was used in previous CBPP studies (Smith et al. 2016; Kashyap et al. 2019) and could be considered as an optimal combination of ridge regression and LASSO, harnessing both the high prediction performance and sparse representation (Zou and Hastie 2005), we here focused on ridge regression and EN.

Psychometric prediction is typically performed by training and testing regression algorithms through cross-validation. For each combination of approaches, we performed 10 repeats of 10-fold cross-validation. As the HCP cohort consists of families of siblings, the cross-validation folds were generated while ensuring that family members were always kept within the same fold, as done in previous studies (Kong et al. 2019; Li et al. 2019). Using Matlab’s regress function, the 9 confounding variables were regressed out from the training set from 9 folds; the same regression coefficients were used to remove effects of these variables from the test set in the remaining fold. Similarly, feature selection for the preliminary evaluation was performed based on the training data only; the same selected features were used in the test fold.

The hyperparameter, epsilon, for linear SVR was set to the default value. As the default value is dependent on data variance and usually found to be the most optimal value during our preliminary analysis, our final implementation of linear SVR did not include hyperparameter tuning. Overall, our implementation of linear SVR can be seen as a simpler linear regression model, in comparison to EN which would rather require hyperparameter tuning. Hyperparameter tuning for EN was done in two steps. During each cross-validation run, we first fixed the alpha value, which determines the compromise between ridge and lasso, while tuning the lambda value, which determines the degree of regularization. To do so, a 10-fold inner-loop cross-validation was carried out using 8 of the training folds. The best alpha value was then chosen according to the prediction performance on the last training fold. For RR, the ridge coefficient was chosen using a similar procedure for EN, by designating one training fold to be the inner validation fold. The coefficient value giving rise to the best prediction performance on the inner validation fold was chosen.

### 2.4. Region-based CBPP

To directly investigate the neurobiological validity and hence interpretability of the brain-behavior patterns, we propose a region-based (or parcel-wise) CBPP framework. The procedure of region-based psychometric prediction was overall the same as that shown in Figure S1 and Figure 2. The main difference is that the input features become an individual brain region or parcel’s connectivity profile, which is represented by a vector of FC values between the region and all other regions.

For easy interpretation, we focused on combinations using Pearson correlation (see Discussion). Based on the preliminary evaluation, optimal performance could be achieved using any combinations with FIX or FIX+GSR data, SVR or EN, at 300-parcel granularity. Here we focused on the FIX-Pearson-SVR combination at 300-parcel granularity with the ‘standard’ confound controlling approach, to facilitate the assessment of denoising strategies later.

We first present the region-based CBPP from the brain region’s perspective, in which a psychometric prediction profile could be established for each brain region or parcel, consisting of the prediction accuracies of the 40 psychometric variables using the connectivity profile of that parcel. We examined four pairs of parcels at the cortical surfaces that were identified in the surface-based Shaefer atlas, located in the primary visual cortex, the premotor cortex, the supramarginal gyrus and the Broca region. Additionally, we examined parcels in the hippocampus hence using volumetric data and an independent atlas derived from independent volumetric RSFC data (AICHA atlas which consists of 300 cortical parcels and 84 subcortical parcels, see Joliot et al. 2015). We selected well-studied regions from different functional networks such that their profiles can be compared with the brain mapping literature. This additional analysis allows the comparison of the psychometric profile of the volumetric parcels with their behavioral profile based on activation data (which typically exists in volumetric space).

We then present the psychometric variable’s perspective, in which the prediction accuracies distribution across parcels could be visualized for each psychometric variable. We selected variables that are assumed to reflect overall crystallized cognition and an intensively study specific domain of cognition (working memory). For comparison, we included one variable pertaining to a further specific paradigm condition within that domain (working memory for faces). Finally, we added one variable related to motor functions (strength).

Finally, to illustrate the issue of interpretation by relating connectivity feature relevance to weights assigned by regression algorithms, we obtained the weight assignments for the same psychometric variables used in illustrating the psychometric variable’s perspective. To simulate a global approach, weights were retrieved for all region-to-region connectivity edges using whole-brain CBPP used in our preliminary evaluation. For each parcel, its regression weight can be represented by the average weight assigned across all its connection with other parcels. Essentially, a highly relevant connectivity edge would increase the regression weights of both parcels in the connection. These regression weights were further averaged across a 10-fold cross-validation loop. The regression weights distributions can then be compared to the prediction accuracies distributions.

For region-based CBPP, no feature selection needs to be done since the number of features is low. For establishing the psychometric profiles, permutation testing was performed by repeating 10-fold cross-validation 1000 times while shuffling the psychometric variables. The final prediction accuracies across the 40 psychometric variables were corrected for multiple comparisons using false discovery rate (FDR; Benjamini & Hochberg 1995) of q < 0.05. For demonstrating the prediction accuracies distribution across parcels, the same permutation testing results were used for assessing statistical significance, corrected for multiple comparisons across parcels.

### 2.5. Effects of denoising and confounds

A potential application of the region-based CBPP framework is to explore the influence of specific parameters in the prediction procedure on the validity of the results. Here we focus on investigating the effects of denoising and confound removal.

To explore the influence of denoising on the validity of the brain-behavior associations, we compared parcel-wise prediction accuracy between minimal preprocessing and FIX denoising data. The overall accuracy difference could be visually discerned, while the difference in variance of prediction accuracies was tested using the Levene’s test (Levene 1960). To further investigate if FIX denoising contributes to the differentiation between parcels’ psychometric profiles or psychometric tests’ prediction accuracy distribution maps, we computed the Euclidean distance between 1) the psychometric profiles and 2) accuracy distribution maps for minimally processed data and for FIX data separately.

To investigate the impact of confounds, we implemented the combination of FIX data followed by SVR using ‘no confound’ approach and ‘sex + brain size confounds’. In the ‘no confound’ approach, the confound controlling step was completely skipped. In the preliminary analysis, when refraining from confounds removal, we found strikingly high prediction accuracy for strength, with further evidence that sex, brain size and intracranial volume were highly correlated with strength (Pearson r = 0.77, 0.54, 0.51 respectively). To further investigate whether these confounds account for the inflated prediction performance, in the ‘sex + brain size confounds’ approach, only these confounds were regressed out.

## 3. Results

### 3.1. Preliminary evaluation

Figure 3 and S3 show the whole-brain CBPP results from all 92 different combinations of approaches. Each point on the line plot shows the average prediction accuracy (Pearson correlation and inversed nRMSD respectively) across the 40 psychometric variables, for a specific combination of approaches. Numerically, the highest average correlation accuracy was achieved by the minimal-partial-SVR combination at 300-parcel granularity, followed by FIX+GSR-partial-SVR and FIX-partial-SVR combination at 300-parcel granularities. The highest average nRMSD accuracy was achieved by FIX-Pearson-EN and FIX+GSR-Pearson-EN combinations. More detailed comparison of approaches including statistical test results can be found in Supplemental materials.

**Figure 3.**
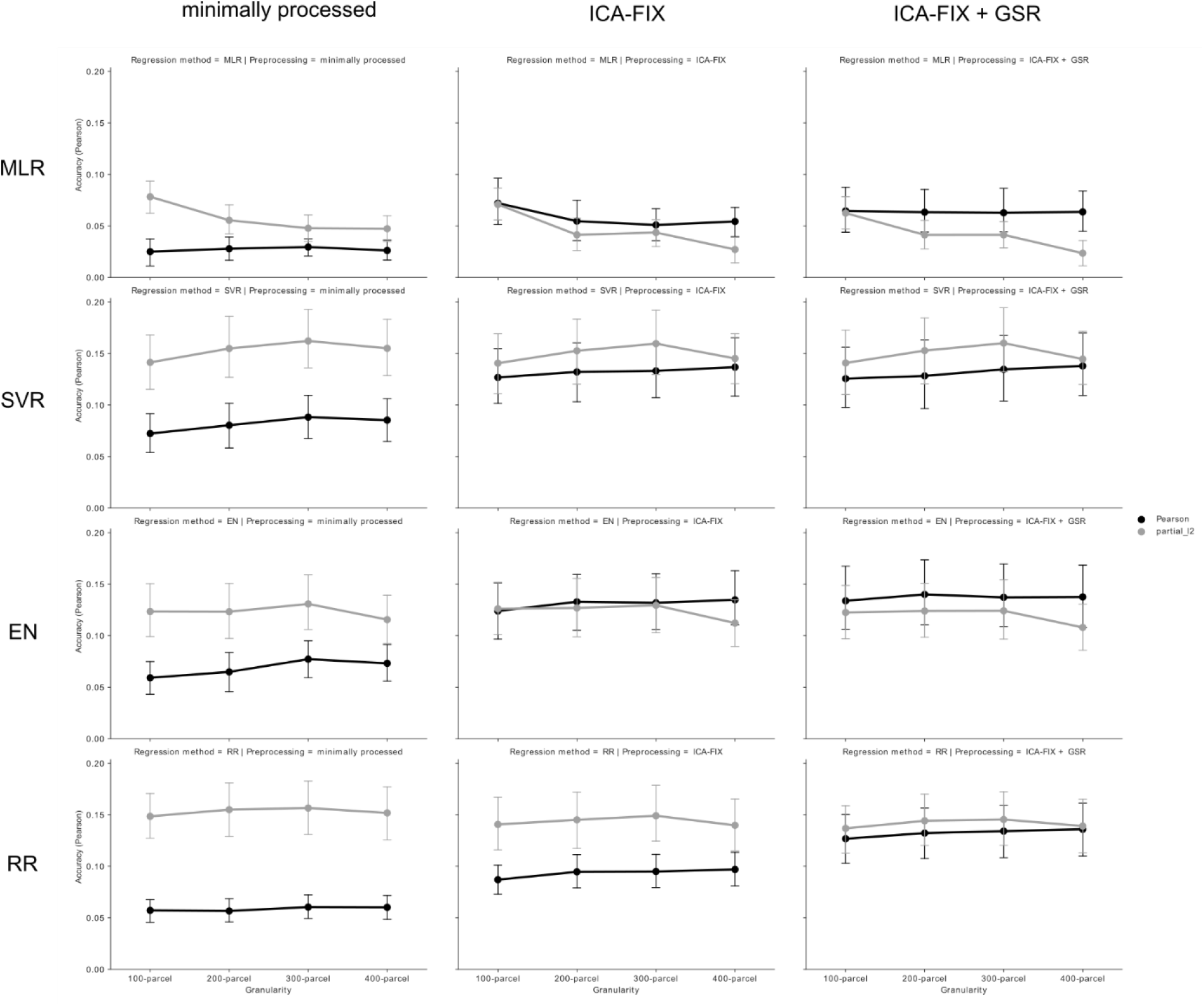
Average prediction accuracy (Pearson correlation between predicted and observed values) across the 40 psychometric variables for each combination of approaches in whole-brain CBPP. Error bars represent the 95% confidence interval across psychometric variables. The columns show combinations using minimally processed, FIX and FIX+GSR data respectively; the rows show combinations using MLR, SVR, EN and RR respectively. Dark lines represent combinations using Pearson correlation, while light grey lines represent combinations using partial correlation.

Overall, machine learning-based denoising (FIX) and a parcellation granularity of 300-parcel led to the most optimal combinations. Following FIX or FIX+GSR denoising and 300-parcel granularity, similar prediction performance could be achieved using any connectivity computation method or regression method (except MLR which led to lower performance).

### 3.2. Brain regions’ psychometric profiles

Figures 4 to 7 show the psychometric profiles of 4 pairs of surface parcels, corresponding to parcels in the primary visual cortex, premotor cortex, supramarginal gyrus and Broca region. Overall, the selected parcels showed similar psychometric profiles across hemispheres, but distinct profiles across parcel locations. Psychometric variables describing general cognitive abilities tended to be relatively better predicted across combinations of approaches in whole-brain CBPP and, relatedly, across parcels in parcel-wise CBPP. For example, the total cognition composite score was the fourth best predicted across combinations of approaches on average and among the best predicted variables for all the 4 selected parcels.

**Figure 4.**
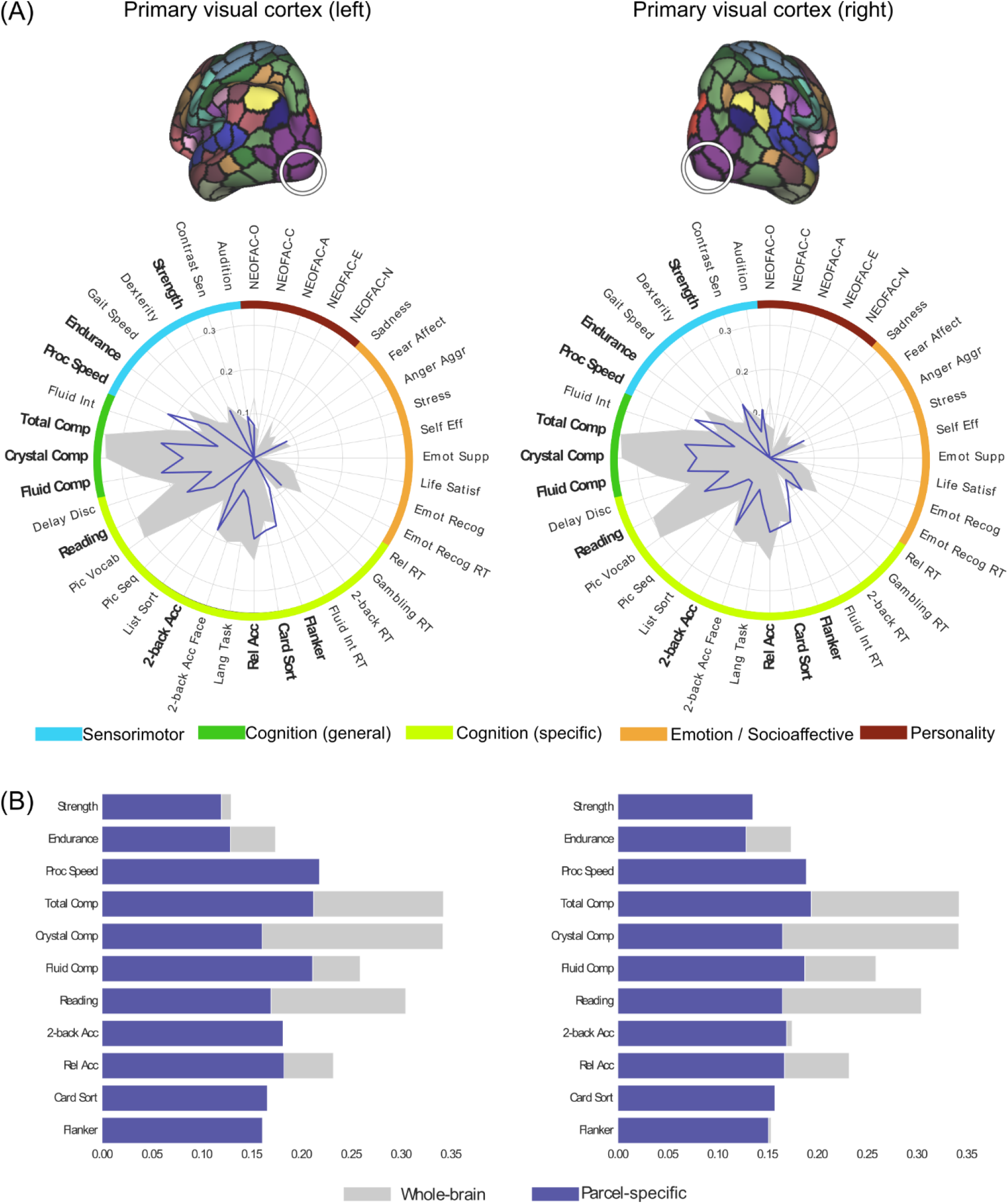
(A) Psychometric profiles for pairs of parcels in primary visual cortex in left and right hemispheres respectively, using FIX-Pearson-SVR combination at 300-parcel granularity, based on Pearson correlation accuracy. Psychometric variables for which the nRMSD accuracy is also significant are highlighted with a bold font. Gray filled contour shows whole-brain prediction profile, while blue contour shows parcel-wise prediction profile. (B) Comparison of parcel-specific (blue) and whole-brain (gray) accuracies for psychometric variables for which both the Pearson and nRMSD accuracies are significant.

**Figure 5.**
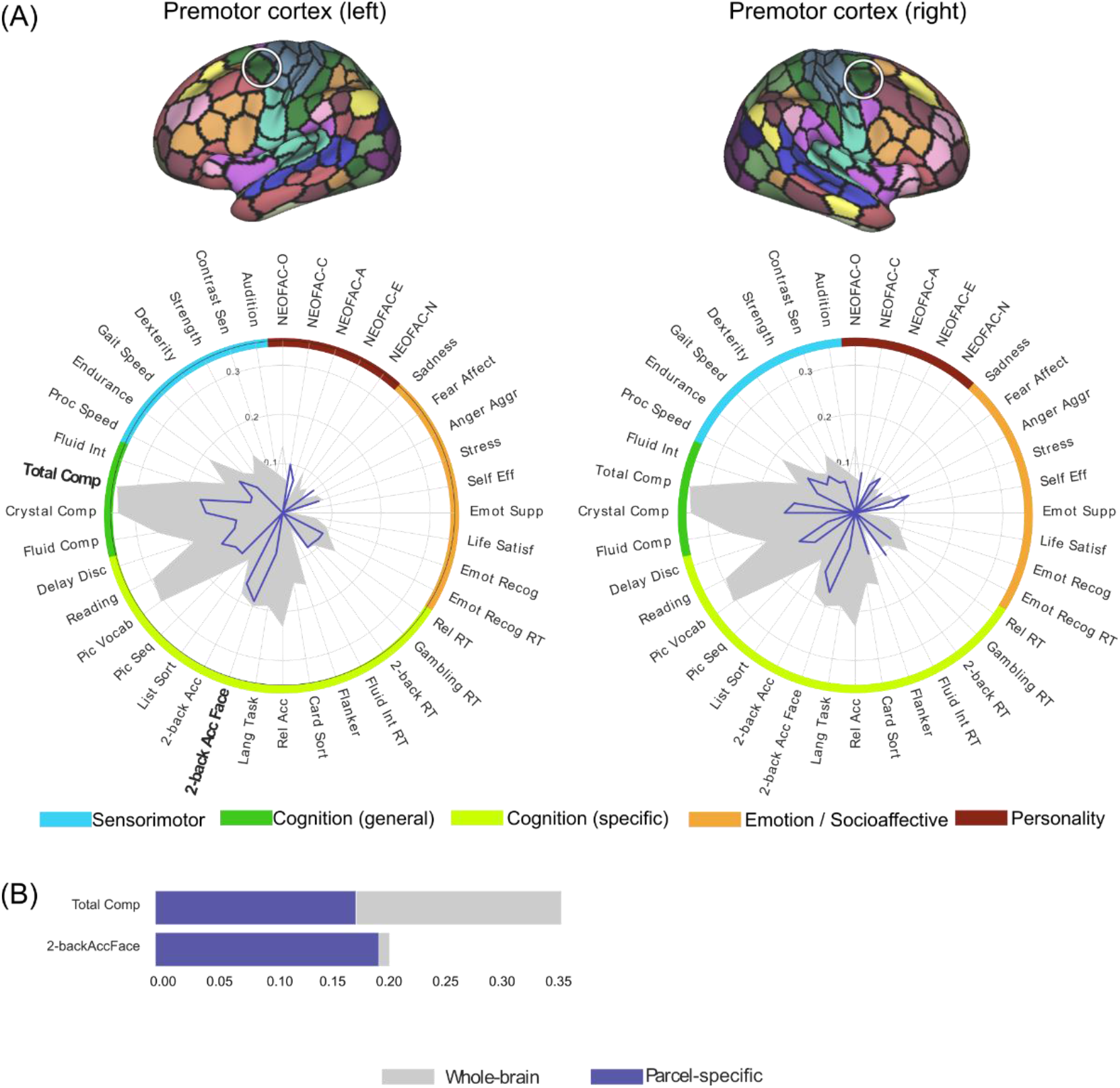
(A) Psychometric profiles for pairs of parcels in premotor cortex in left and right hemispheres respectively, using FIX-Pearson-SVR combination at 300-parcel granularity, based on Pearson correlation accuracy. Psychometric variables for which the nRMSD accuracy is also significant are highlighted with a bold font. Gray filled contour shows whole-brain prediction profile, while blue contour shows parcel-wise prediction profile. (B) Comparison of parcel-specific (blue) and whole-brain (gray) accuracies for psychometric variables for which both the Pearson and nRMSD accuracies are significant.

**Figure 6.**
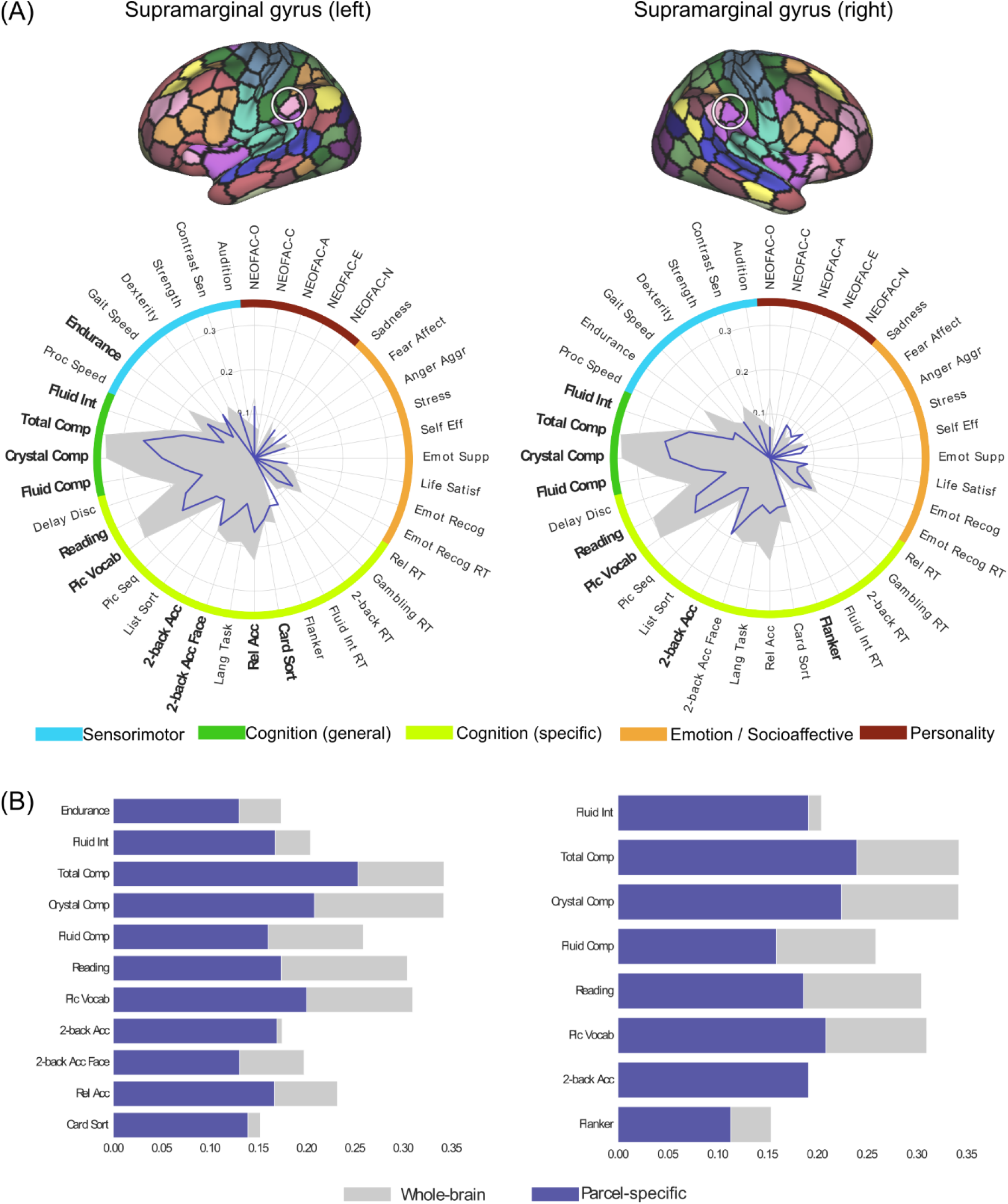
Psychometric profiles for the pair of parcels in supramarginal gyrus in left and right hemispheres respectively, using FIX-Pearson-SVR combination at 300-parcel granularity, based on Pearson correlation accuracy. Psychometric variables for which the nRMSD accuracy is also significant are highlighted with a bold font. Gray filled contour shows whole-brain prediction profile, while blue contour shows parcel-wise prediction profile. (B) Comparison of parcel-specific (blue) and whole-brain (gray) accuracies for psychometric variables for which both the Pearson and nRMSD accuracies are significant.

**Figure 7.**
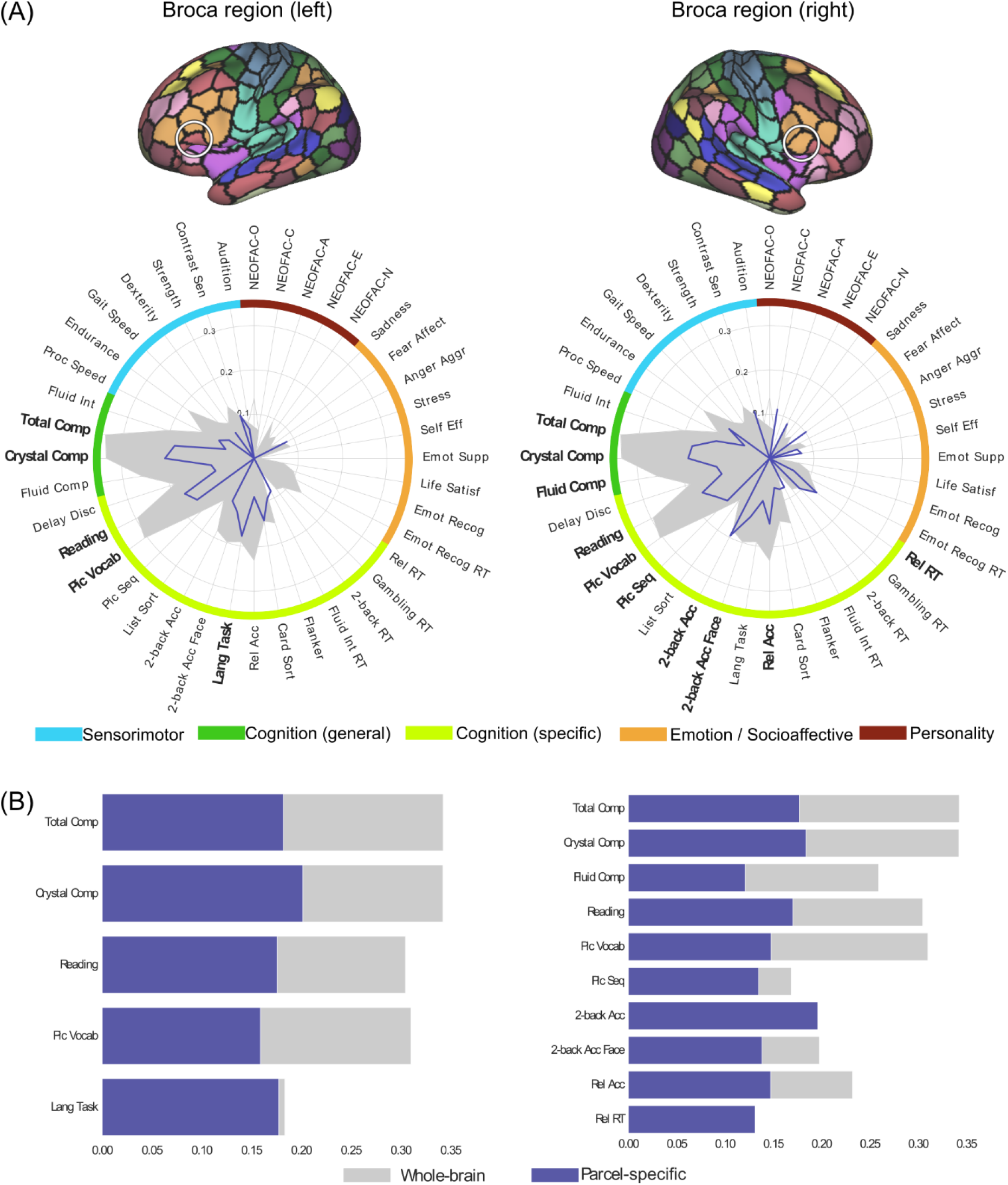
Psychometric profiles for the pair of parcels in Broca region in left and right hemispheres respectively, using FIX-Pearson-SVR combination at 300-parcel granularity, based on Pearson correlation accuracy. Psychometric variables for which the nRMSD accuracy is also significant are highlighted with a bold font. Gray filled contour shows whole-brain prediction profile, while blue contour shows parcel-wise prediction profile. (B) Comparison of parcel-specific (blue) and whole-brain (gray) accuracies for psychometric variables for which both the Pearson and nRMSD accuracies are significant.

We note that the two prediction performance measures, Pearson correlation and inversed nRMSD, converge on highlighting similar variables. However, discrepancies can also be observed as the nRMSD measure appears to be much stricter than the correlation measure. Therefore, we suggest to focus on patterns for which convergence across measures are observed as these might be more reliable than any pattern yielded by a single prediction performance measure.

When analyzing the brain regions’ psychometric profiles, we focus on the relative comparisons of prediction accuracy between different psychometric variables, for each parcel separately. While most of the relatively better predicted psychometric variables for parcels in the primary visual cortex were also relatively better predicted by the connectivity profiles of many other brain regions, processing speed (*Proc Speed*) was predicted with relatively better accuracy only in the primary visual cortex parcels. In contrast, FC patterns of the primary visual cortex parcels showed nearly no predictive power for psychometric variables in the emotion domain. In comparison to other parcels, the premotor cortex parcels showed overall low prediction accuracies for most psychometric variables. The parcels in the supramarginal gyrus showed relatively better predictive power in most cognition-related variables, including total cognition composite score, fluid intelligence (*Fluid Int*), working memory performance, picture vocabulary (*Pic Vocab*) and crystallized cognition composite score (*Crystal Comp*), in comparison to variables from other domains. Finally, the parcels in the Broca region showed relatively better prediction power for language-related measures, cognition composite scores and working memory performance than for other psychometric variables. Interestingly, the right parcel showed worse prediction power for language task accuracy (*Lang task*) than the left, but better prediction power for working memory abilities than the left. Figure 8 shows the psychometric profiles of a pair of parcels in the anterior hippocampus. Across hemispheres, the two parcels showed similar psychometric profiles. Again, both parcels showed relatively better predictive power for all cognition composite scores, processing speed, reading and picture vocabulary, cognitive flexibility (*Card Sort*) and endurance than for other psychometric variables. In addition, the left anterior hippocampus showed relatively better prediction accuracies in extraversion (*NEOFAC-E*), while the right parcel showed relatively better prediction accuracies in relational task performance (*Rel Acc*), than in other psychometric variables. A behavioral profiling of the hippocampus parcels across activation studies of the BrainMap database (Fox and Lancaster 2002; Laird et al. 2005), shown in figure S4, revealed that these parcels are consistently activated for emotion, memory, face processing and passive viewing paradigms, hence partly converging with their psychometric profile based on CBPP.

**Figure 8.**
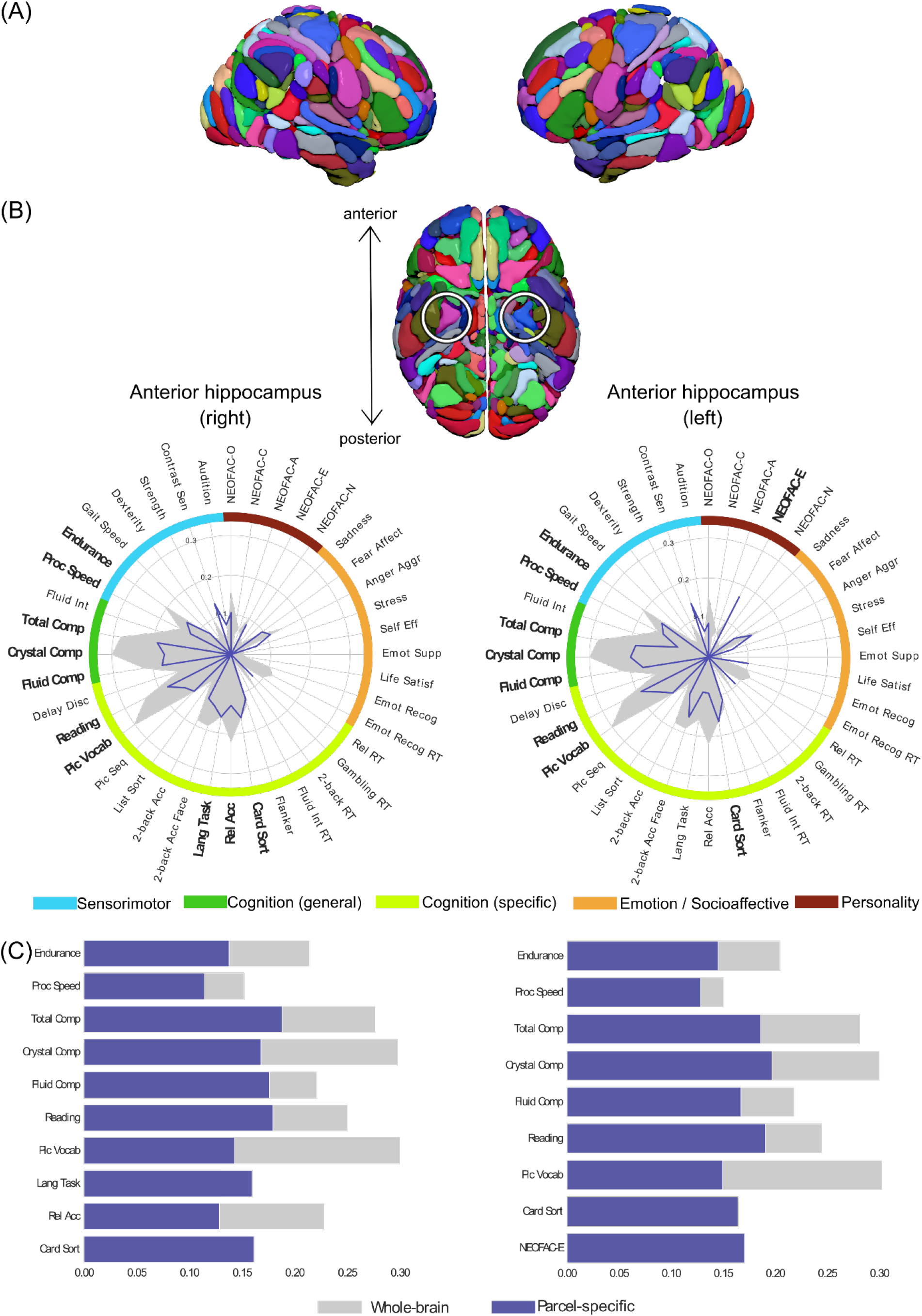
(A) The AICHA atlas (B) Psychometric profiles for the pair of parcels in anterior hippocampus in right (pink parcel) and left (blue parcel) hemispheres respectively, using FIX+WM/CSF-Pearson-SVR combination, based on Pearson correlation accuracy. Psychometric variables for which the nRMSD accuracy is also significant are highlighted with a bold font. Gray filled contour shows whole-brain prediction profile, while blue contour shows parcel-wise prediction profile. (B) Comparison of parcel-specific (blue) and whole-brain (gray) accuracies for psychometric variables for which both the Pearson and nRMSD accuracies are significant.

Panel B in figures 4 to 8 shows the comparison of parcel-specific and whole-brain prediction accuracy, for psychometric variables where both the Pearson correlation and nRMSD measures showed statistical significance. For almost all parcels, the parcel-wise accuracies for the 3 cognition composite scores were about half as much as the corresponding whole-brain accuracies. Similar ratios were also observed for oral reading and picture vocabulary when their parcel-wise accuracies were selected for comparison. On the other hand, several psychometric variables showed at least comparable parcel-wise accuracies to the whole-brain accuracies. For the parcels in the primary visual cortex, such variables include processing speed, working memory performance, cognitive flexibility and inhibitory control (*Flanker*). For the parcels in the premotor cortex, these include working memory performance for face material for the left hemisphere. For the parcels in the supramarginal gyrus, these include working memory performance for both hemispheres, as well as cognitive flexibility for the left hemisphere. For the parcels in the Broca region, these include language task for the left hemisphere, as well as working memory for the right hemisphere. For the parcels in the anterior hippocampus, these include cognitive flexibility for both hemispheres and extraversion for the left hemisphere.

### 3.3. Psychometric variables’ prediction accuracy distributions

From the psychometric variables’ perspective, we present the prediction accuracies distribution across parcels. Figure 9a to 9d show the prediction accuracy distributions and histograms across the brain for 4 selected psychometric variables, using FIX-Pearson-SVR combination at 300-parcel granularity based on Pearson correlation accuracy. Figure S5 shows the prediction accuracy and histograms for the same variables and combinations based on nRMSD accuracy. Across the brain, prediction accuracies were generally low for strength.

**Figure 9.**
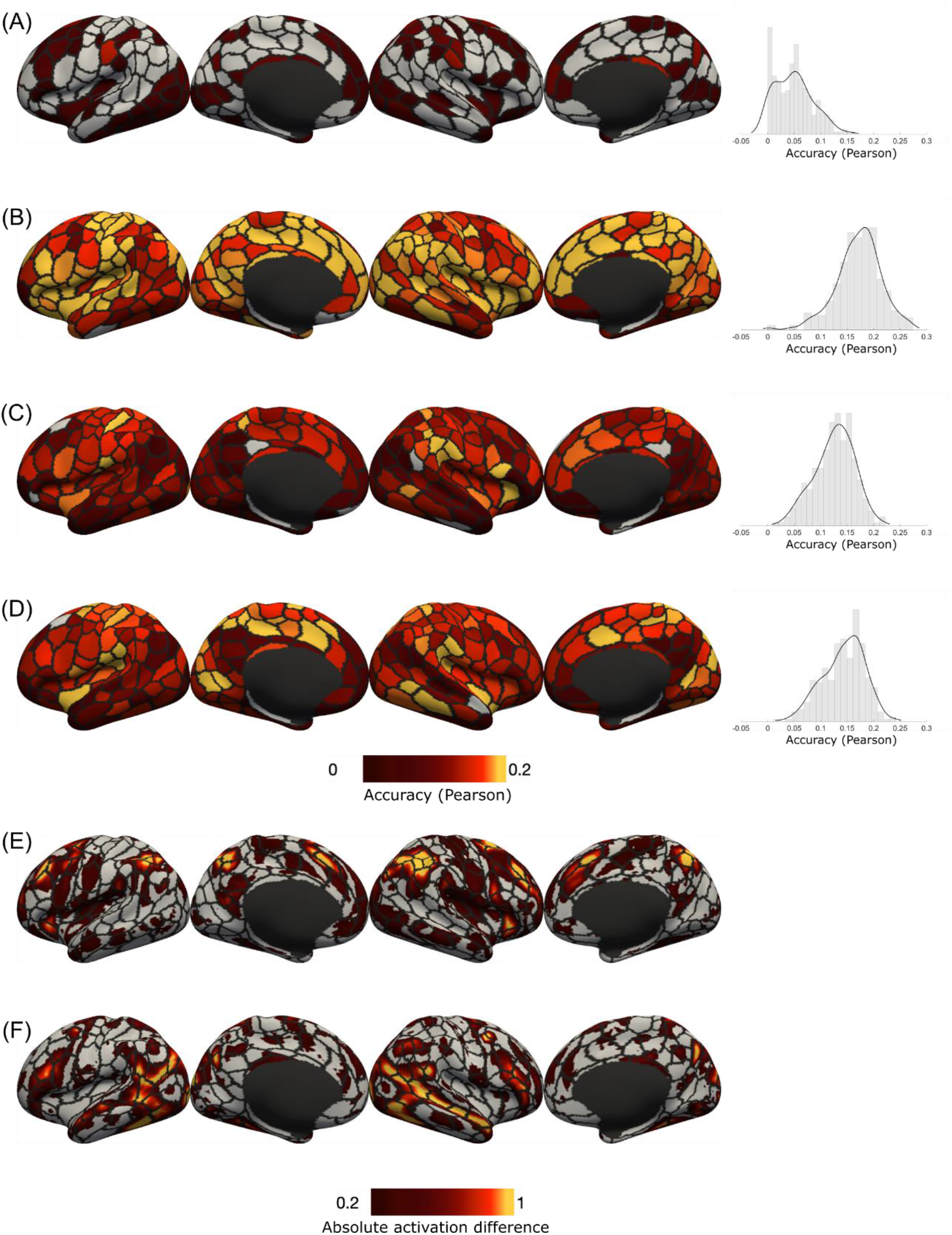
**Top**: Prediction accuracy distribution using FIX-Pearson-SVR combination at 300-parcel granularity of the 4 selected psychometric variables: (a) strength (Strength) (b) crystallized cognition composite score (Crystal Comp) (c) working memory task overall accuracy (2-back Acc) (d) working memory task face condition accuracy (2-back Acc Face). Each row shows the distribution overlayed on the fsLR surface in lateral and medial views of the left and right hemispheres, respectively. Color represents the magnitude of the Pearson correlation coefficients between predicted and observed values. Accuracies which were not statistically significant are shown in gray. The rightmost column shows the histogram of prediction accuracies for each variable respectively. **Bottom**: Absolute values of HCP group activation maps of (e) working memory task (2-back – 0-back) (f) social task (theory of mind – random), with overlay of Schaefer atlas at 300-parcel granularity. Each row shows the activation map overlayed on the fsLR surface in lateral and medial views of the left and right hemispheres, respectively. Color represents the absolute value of the activation difference in the maps. The maps were thresholded to show value in the range between 0.2 and 1 (roughly the 50^th^ and 99^th^ percentile respectively). Vertices outside the threshold range were shown in gray.

When analyzing the psychometric variables’ prediction accuracy distributions, we focus on the relative comparisons of prediction accuracy between different parcels, for each psychometric variable separately. For strength prediction, the better performing parcel was the parcel in the ventral postcentral sulcus in the left hemisphere. For crystallized cognition composite score, many parcels achieved similar performance. For working memory task overall accuracy (*2-back Acc*), the better performing parcels were those in the cingulate cortex, parietal cortex, supramarginal gyrus, lateral frontal cortex and anterior insula. For working memory task performance specifically for faces (*2-back Acc Face*), the distribution was similar to that of the working memory task overall accuracy; in addition, parcels in the inferior temporal cortex and the calcarine sulcus achieved similar performance in predicting working memory abilities for faces.

Figure 9e and 9f shows the HCP group activation maps for working memory task (*tfMRI_WM_2BK-0BK*) and social task (*tfmri_SOCIAL_TOM-RANDOM*), where activations are shown in absolute values. Overall, the pattern in the prediction accuracy distribution for working memory overall accuracy is more similar to the working memory activation pattern, than with the social task activation pattern. In particular, high signal changes in working memory task and high prediction accuracies were both found in the anterior cingulate cortex, parietal cortex, lateral frontal cortex and anterior insula. In contrast, high signal changes in social task were found across the ventral network including the temporal lobe, where prediction accuracies were generally low for working memory performance.

Figures S6 to S9 shows the Pearson correlation accuracy distribution using FIX-Pearson-SVR combinations across all 4 granularities, for the 4 selected psychometric variables respectively. By visual inspection, the overall distribution patterns were similar across different granularities. Nevertheless, within broad functional territories, the relatively better predictive power of specific parcels (hence subregions), in comparison with the rest of the parcels, appeared when reaching granularities such as 200-parcels and 300-parcels, which could reflect relatively more specific brain region-behavior relationships captured at higher granularities.

### 3.4. Prediction accuracy distribution vs. regression weights

Figure 10a to 10d shows the absolute mean (top rows) of each parcel’s regression weights for the 4 selected psychometric variables across cross-validation. In comparison to the prediction accuracy distributions (Figure 9), the only obvious similarity is the high relevance of a left middle frontal parcel and some right cingulate parcels for crystallized cognition composite score. While it might be possible to find a number of connections with high regression weights assigned across cross-validation, it was often not possible to group them in a meaningful manner. As illustrated in Figure 10, the average regression weights for a parcel is at most about 0.04 (for a whole-brain prediction model that itself explains only 1% to 10% of the variance in the psychometric variables). Consequently, it would not be sound to claim that any parcel was found to be importantly related to the predicted psychometric variable. Furthermore, the distribution of mean regression weights suffers from a lack of smoothness and hemispheric similarity, questioning its neurobiological validity. In particular, the Pearson correlations of regression weights across parcels between hemispheres were not statistically significant (r = 0.06, 0.02, -0.01 and 0.02, p = 0.43, 0.81, 0.94 and 0.83 for the four psychometric variables respectively; Student’s t distribution with alpha < 0.05). On the other hand, the Pearson correlations of region-wise prediction accuracies between hemispheres were all significant (r = 0.25, 0.20, 0.34 and 0.48 for the four psychometric variables respectively, all p < 0.01; Student’s t distribution with alpha < 0.05).

**Figure 10.**
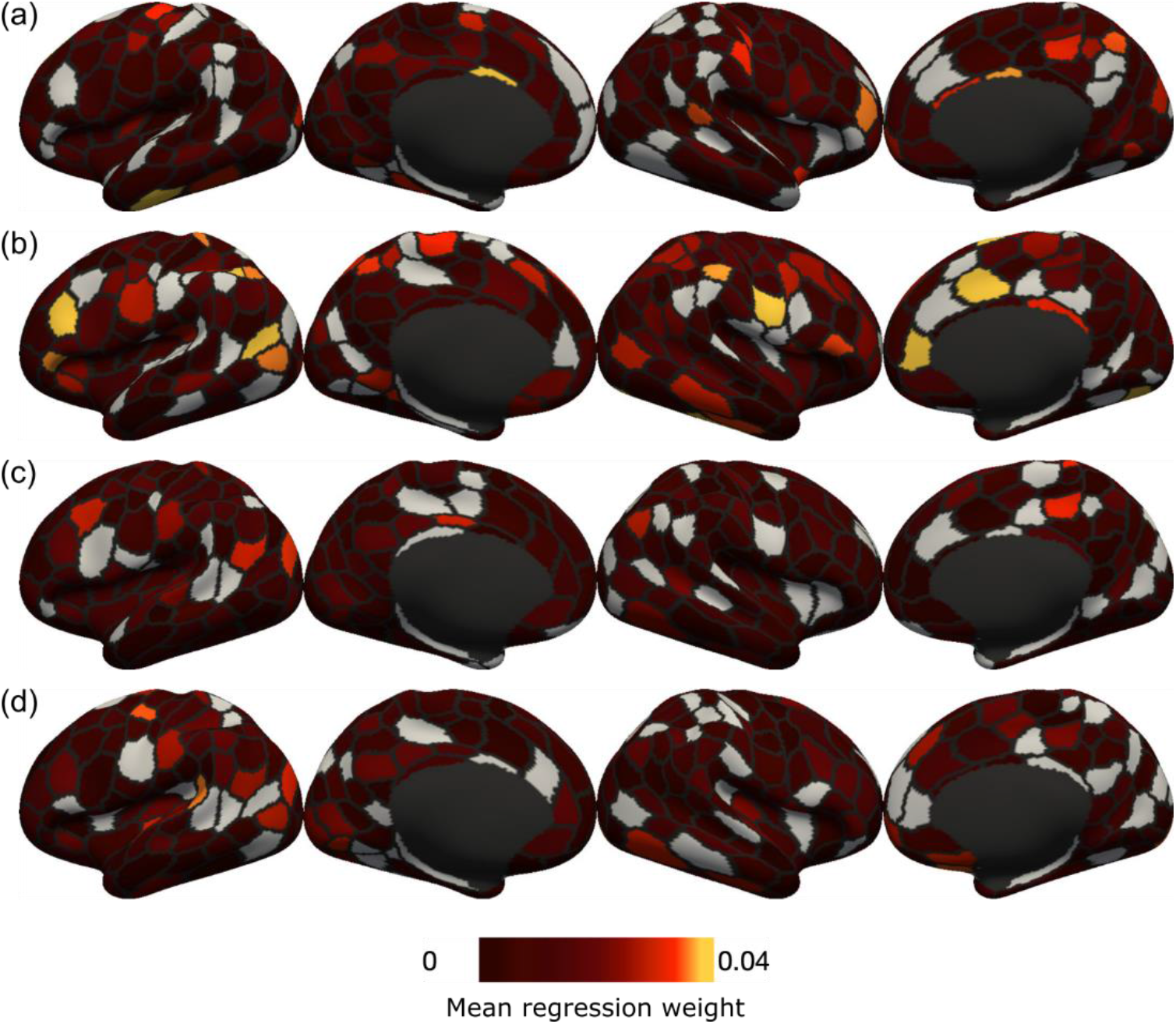
Assessing parcels’ relevance by regression weights using FIX-Pearson-SVR combination at 300-parcel granularity for the 4 selected psychometric variables: (a) strength (Strength) (b) crystallized cognition composite score (Crystal Comp) (c) working memory task overall accuracy (2-back Acc) (d) working memory task face condition accuracy (2-back Acc Face). Each row shows the activation map overlayed on the fsLR surface in lateral and medial views of the left and right hemispheres, respectively. Color represents the average weight value assigned across cross-validation. Values close to zero were shown in gray.

### 3.5. Effects of denoising

Figure 11 shows the prediction accuracy distribution and histogram of parcel-wise CBPP across the brain for 4 selected psychometric variables, using minimal-Pearson-SVR combination at 300-parcel granularity, based on Pearson correlation accuracy. Similarly, figure S10 shows the prediction accuracy distribution and histograms based on nRMSD accuracy. Overall, the prediction accuracies were lower compared to those using FIX data. Furthermore, the variance of prediction accuracies across parcels were significantly lower for minimally processed data than for FIX data, for the strength variable (Levene’s test p < 0.01 for both Pearson and nRMSD accuracies; Levene 1960), suggesting that parcels were more differentiated after FIX denoising. For the Euclidean distance comparison, in all cases, the Euclidean distances were significantly larger following FIX denoising than following minimal processing (p < 0.0001), suggesting that both psychometric profiles of different parcels and prediction accuracy distributions for different psychometric variables were more dissimilar when FIX denoising was used.

**Figure 11.**
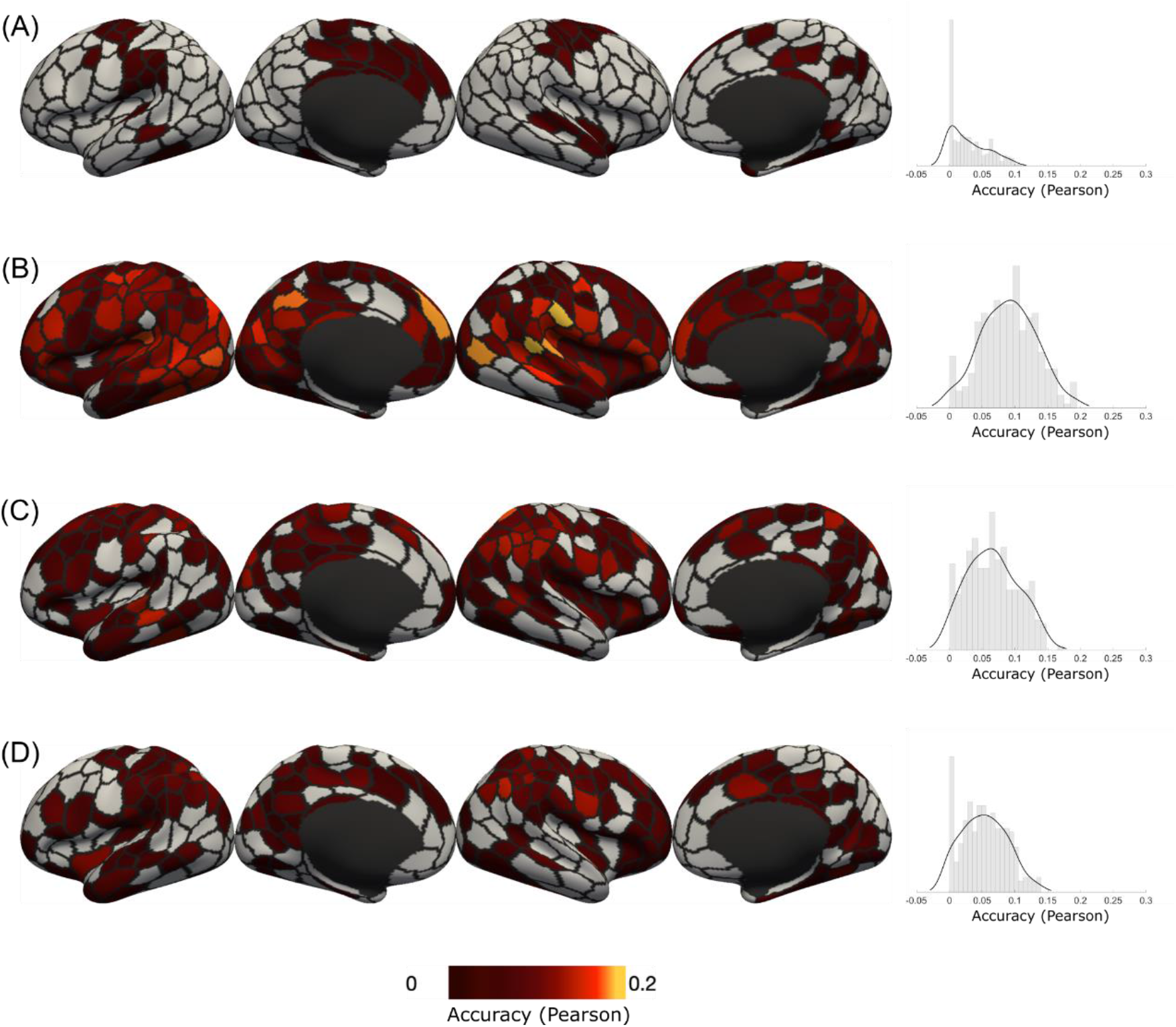
Prediction accuracy distribution using minimal-Pearson-SVR combination at 300-parcel granularity of the 4 selected psychometric variables: (a) strength (Strength) (b) crystallized cognition composite score (Crystal Comp) (c) working memory task overall accuracy (2-back Acc) (d) working memory task face condition accuracy (2-back Acc Face). Each row shows the activation map overlayed on the fsLR surface in lateral and medial views of the left and right hemispheres, respectively. Color represents the magnitude of the Pearson correlation coefficients between predicted and observed values. Accuracies which were not statistically significant are shown in gray. The rightmost column shows the histogram of prediction accuracies for each variable respectively.

### 3.6. Effects of confounds

Figure 12 shows the ‘no confound’ and ‘sex + brain size confounds’ prediction results for pairs of parcels in the supramarginal gyrus and Broca region respectively. When the ‘no confound’ approach was used, extremely high prediction accuracies for strength were observed, even though both regions are mainly cognition-related. This suggests that not controlling for confounding variables (such as brain size) could undermine the interpretability of parcel-wise CBPP. Our standard regression approach appeared to greatly neutralize such aberrant prediction performance, while simply regressing out sex and brain volume-related confounds had a similar effect.

**Figure 12.**
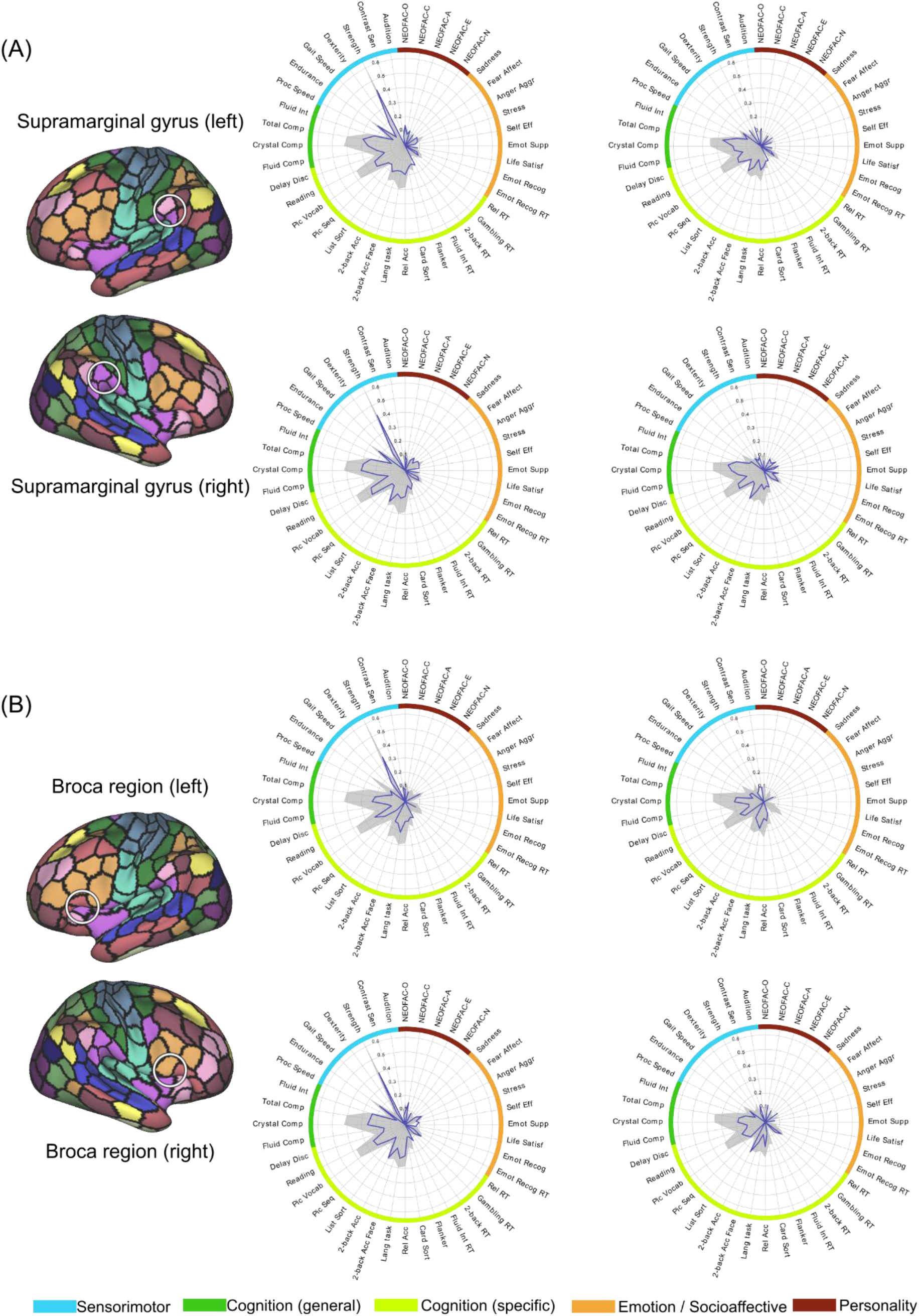
Psychometric profiles for pairs of parcels in (a) Supramarginal gyrus (b) Broca region in left and right hemispheres respectively, using FIX-Pearson-SVR combination at 300-parcel granularity with ‘no confound’ (left) and ‘sex + brain size confounds’ (right) approach. Gray filled contour shows whole-brain prediction profile, while blue contour shows parcel-wise prediction profile

## 4. Discussion

To develop an optimal framework of connectivity-based prediction of psychometric variables (CBPP) for cognitive neuroscience, we first evaluated the effects of different approaches and parameters that have been used in whole brain or network-based approach in previous studies. Overall, our results demonstrated the relevance of sophisticated denoising approaches for resting fMRI data, as well as the good performance of standard regression-based prediction algorithms. Capitalizing on this preliminary investigation, we then demonstrate the validity of our region-wise CBPP approach, in terms of single brain region’s psychometric profile as well as single psychometric variable’s prediction accuracy distribution.

Our findings show the benefit of sophisticated machine learning-based denoising (here FIX denoising), not only at the quantitative level on prediction performance but also at the qualitative level for the study of brain-behavior relationships. We can note however that, while prediction accuracies were significantly lower for minimally processed data in comparison to FIX and FIX+GSR data when Pearson correlation was used, the three different preprocessing approaches were comparable when partial correlation was used. Considering that the computation of partial correlation between two parcels, to a certain extent, implies the neutralization of common variance, it is possible that using partial correlation potentially removes noise and artifacts affecting several parcels. As a result, similar denoising effects could be achieved by computing partial correlation and by FIX (or FIX+GSR) preprocessing (also see figure S11). Nevertheless, our supplementary investigation (see figure S11) suggests that partial correlation aggressively alters the complex pattern of connectivity fingerprint of brain region (i.e. parcels) by removing shared variance across regions. In line with this view, our preliminary analysis (see figure S12) revealed that the residual connectivity fingerprint of the region following partial correlation does not lead to distinguishable psychometric profiles for different parcels. Consequently, these combinations were excluded from the final analysis. At the current stage of knowledge, we can only assume that the common variance removed in partial correlation computation contains variance of interest (e.g. common variance of parcels in the same functional network) that importantly contributes to the region’s FC fingerprint. Therefore, while using partial correlation may improve overall prediction performance, it may not provide the same level of interpretability as models using Pearson correlation gives when using region-wise predictions.

Our qualitative examination of the psychometric prediction profile for each parcel generally matched well with our expectations based on brain mapping literature. Hence, the parcels in primary visual cortex showed predictive power for processing speed performance. This pattern could be explained by the role of these regions in visual information processing and relaying (Fabre-Thorpe et al. 2001; Sharpee et al. 2006). The performance at two higher cognitive variables (executive function type: *Card Sort* and *Flanker*) were also relatively better predicted from primary visual cortex’s FC patterns, which could potentially be explained by the role of visual processing speed and accuracy in the performance. The parcel in left premotor cortex showed predictive power for working memory performance for face material, which is in line with the premotor cortex’s engagement in working memory and face processing paradigms (Chan and Downing 2011; Balconi and Bortolotti 2013; Genon et al. 2017, 2018). The parcels in supramarginal gyrus showed predictive power across a wide range of higher cognitive function measures. Additionally, overall, the FC patterns of the parcels in supramarginal gyrus and the Broca region show predictive power for psychometric variables related to language processing and crystallized intelligence. Both regions have been previously shown to be involved in word and phonological processing (Binder et al. 1997; McNealy et al. 2006; Klepousniotou et al. 2014; Twomey et al. 2015; Oberhuber et al. 2016). Comparing the prediction profile of the left and right Broca region further supports the validity of our approach since, in contrast to the left Broca region, the right Broca region showed relatively low predictive power for language task performance, a pattern that is in agreement with the left hemisphere’s dominance for language function (Clos et al. 2013; Karsolis et al. 2019). Furthermore, parcels in these two regions also showed predictive power for working memory measures, in agreement with their role in the phonological loop in traditional working memory models (Salmon et al. 1996; Smith and Jonides 1999; Zurowski et al. 2002; Rogalsky and Hickok 2011). Finally, the anterior hippocampus also shows some predictive power for general cognition performance, including reading and cognitive flexibility. However, differences between hemispheres in psychometric profile were also observed for the hippocampus. Hence, the right hippocampus showed relatively high predictive power for language performance, relational task performance as well as picture vocabulary, while the left hippocampus showed relatively high predictive power for endurance and extraversion. Because the hippocampus parcels were in a volumetric format, their psychometric prediction profile can be compared with behavioral profiling derived from the aggregation of activation data (which is typically in volumetric format). Here, computing the behavioral profile of the anterior hippocampi confirmed that these regions are usually associated with memory and emotion (Moser and Moser 1998; Prince et al. 2005; Plachti et al. 2019). Hence, the psychometric profile of brain regions revealed here partly converge with the a-priori information from the human brain mapping literature, including when a quantitative statistical approach is used to summarize this literature (but see last paragraph for a discussion on the limitations).

Across the parcels, the comparisons between parcel-wise and whole-brain prediction accuracies revealed two different trends. For multidetermined psychometric variables like the cognition composite scores, while most parcel-wise predictions were statistically significant, their accuracies usually amounted to half the whole-brain accuracies. Such findings reinforced the conceptual view according to which such general cognitive performance is supported by wide-spread distributed networks rather than individual regions. On the other hand, performance at several psychometric variables could be predicted roughly equally well by the region-based and the whole-brain approach, like cognitive flexibility and working memory performance. In these cases, it is possible that the connectivity profile of some specific regions would contain crucial information for the prediction, so that the whole-brain connectivity matrix does not necessarily convey more relevant signal than the parcel-wise connectivity features from a predictive standpoint. Future studies could further investigate if region-based approach could predict some psychometric variables better than the whole-brain approach and the implications, both from a conceptual cognitive neuroscience point of view and from an applied perspective. In the current work, we focus on optimizing our region-based approach and validating the additional insights provided by combining it with whole-brain approaches. Neurobiological validation was assessed by showing the convergence with the brain mapping literature. We would here like to emphasize that, to the best of our knowledge, the neurobiological validity or utility for cognitive neuroscience of predictive models is very rarely investigated in connectivity-based studies. Most studies adopted an a-priori focus on specific networks or regions to build a predictive model and are, in that sense, not truly data-driven, while other studies derived neuroscientific interpretations from the weights assigned to the features, an approach which has been shown to be misleading (Haufe et al. 2014). Thus, we believe that our study addresses an important gap in the current brain-based prediction literature.

In the same vein, our examination of the psychometric prediction accuracy spatial distribution maps appeared to converge also well with the brain mapping literature. When examining specific measures such as strength performance, our prediction maps revealed a relatively specific pattern reminiscent of the sensorimotor network. In contrast, multidetermined score such as the crystallized intelligence composite score revealed a very broad pattern, in which the parcels with the best prediction power lie in regions that frequently show high interindividual variability in RSFC patterns, such as the supramarginal gyrus and the anterior insula (Mueller et al., 2013; Laumann et al., 2015; Kong et al., 2019). Comparing the pattern of prediction performance for the total accuracy score for the 2-back task and the same score only for faces material revealed overall similar patterns but with better prediction power in the right hemisphere for faces than for the general 2-back task, in particular in the ventral temporal regions, in agreement with the literature on faces processing in the brain (Sams et al., 1997; Nakamura et al., 2000; Nelson, 2001).

When examining the accuracy distribution maps across granularities level (i.e. region subdivision level), the overall patterns remain similar for every selected psychometric measure. With increased granularity, especially at 200-parcel and 300-parcel granularity, the proportion of parcels showing better predictive power compared to the proportion of parcels showing genuinely low predictive power increased. This coincides with our observations of achieving better prediction performance in whole-brain CBPP with higher granularity, thus reinforcing the conclusion that 200 to 300 parcels represents an optimal range for fMRI surface data. Finally, our observation that the subdivision of broad functional region into specific parcels at higher granularity goes along with the better predictive power for some parcels (in comparison to the rest of the parcels) may reflect the fact that these parcels represent functional sub-units, and hence that 200/300 parcels granularity provides a better representation of brain functional data from a cognitive neuroscience standpoint.

To sum up, overall, several lines of evidence support the validity of a region-wise connectivity-based psychometric prediction framework. In the absence of ground truth, one line of validation of the brain-behavior pattern here was the comparison with patterns yielded by activation studies. However, comparisons between the two approaches remain conceptually questionable because brain patterns of activation studies typically reflect regions that are consistently activated across samples of participants and thus, regions whose activations should strictly not depend on interindividual variability in the completion of the task (see Genon et al. 2018 for a broader discussion). In contrast, a connectivity-based prediction approach of psychometric data typically capitalizes on the interindividual variability in brain and behavior. Therefore, the question remains fully open on the extent to which both approaches should converge. Our framework could bring further insight into this question in the future. As an additional challenge for the future, our region-wise approach remains limited by some general issues that affects the whole field of connectivity-based psychometric prediction. Namely, the prediction performance remains in a relatively low range, reflecting the limited part of variance in psychometric variables in a healthy adult population that can be explained by interindividual variability in resting-state functional connectivity. To this end, appropriate deconfounding and cross-validation procedures need to be adopted, which could in turn lead to lower prediction accuracies as non-neurobiological variance are removed. Consequently, inferences made based on connectivity-based psychometric prediction are inherently weak and remain to be, at least conceptually, replicated. Accordingly, here, we avoid interpreting results based on the absolute accuracy values, but we focused on the relative predictive power across brain regions or psychometric variables. At first sight, the best predicted scores for both region-wise and whole-brain based predictions are very general scores reflecting global cognitive abilities, such as composite scores, crystallized and fluid intelligence, as well as vocabulary level. We can assume that education (both formal with regards to year spent in academia, as well as informal, as determined by the socio-economic background) plays a major role in this interindividual variability, shaping brain functional connectivity across years and, in turn, optimizing performance at standard cognitive tests. This is the reason why we refrained from regressing variance related to education in the present study. However, the current approach offers a framework to progressively disentangle the influence of factor such as formal education on the relationship between region’s connectivity profile and one’s cognitive abilities. Thus, the current framework shows potential to address open questions and issues in the field in the future.

One general conceptual limitation of our region-based framework would be the negligence of the distributed aspect of brain-behavior relationship, where the behavior in consideration arises from wide-spread distributed networks instead of localized regions (Dubois et al. 2018; Genon et al. 2018). This conceptual perspective is usually referred to as “global” (whole-brain connectivity or network-based connectivity) approaches in the prediction of behavior (Kong et al. 2019; Kashyap et al. 2019; Pervaiz et al. 2020). In line with this view, the global approaches usually lead to higher prediction performance for many behvaioral variables. Accordingly, a global approach should be preferred when the overall prediction performance matters, while our region-based approach helps to evaluate the relative contribution across regions for specific cognitive measures. Importantly, both approaches could be used in conjunction for additional insights into the extent at which distributed or localized certain brain-behavior relationships are, as illustrated in our analysis of brain region’s psychometric profiles.

To conclude, by using high quality data, including a broad range of psychometric measures in a healthy adult cohort, we developed a region-wise connectivity based psychometric prediction framework based on supervised learning approaches linking resting-state functional connectivity of brain regions to behavior. To promote the use of our specific region-wise approach in cognitive neuroscience studies, we illustrated two main applications for which we evaluated the brain region-behavior relationships: 1) psychometric profiles of brain regions, and 2) brain maps of prediction accuracies distribution for specific psychometric variables. The material to implement our approach is openly available at https://github.com/inm7/cbpp. To demonstrate the potential contribution of our region-based approach based on its increased transparency, we illustrated how sophisticated denoising can provide a clarified picture of the association between brain and behavior. We also illustrated the spurious effect of confounds such as brain size on the predictive model. Future work should investigate the potential benefits of utilizing individualized parcellations (Kong et al. 2019). The developed framework could also contribute to a better understanding of the relationships between brain regions’ connectivity and behavioral phenotypes in aging and clinical populations. The transferability of our framework in this context should be investigated in future studies.

## Code availability

All codes are publicly available at https://github.com/inm7/cbpp.

## Acknowledgement

This work was supported by the Deutsche Forschungsgemeinschaft (DFG, GE 2835/1–1, EI 816/ 4–1), the Helmholtz Portfolio Theme ‘Supercomputing and Modelling for the Human Brain’ and the European Union’s Horizon 2020 Research and Innovation Programme under Grant Agreement No. 720270 (HBP SGA1) and Grant Agreement No. 785907 (HBP SGA2). Data were provided by the Human Connectome Project, WU-Minn Consortium (Principal Investigators: David Van Essen and Kamil Ugurbil; 1U54MH091657) funded by the 16 NIH Institutes and Centers that support the NIH Blueprint for Neuroscience Research; and by the McDonnell Center for Systems Neuroscience at Washington University. The authors also thank Hesheng Liu for helpful discussion.

