## Supplemental materials for "A Connectivity-based Psychometric Prediction Framework for Brain-behavior Relationship Studies"

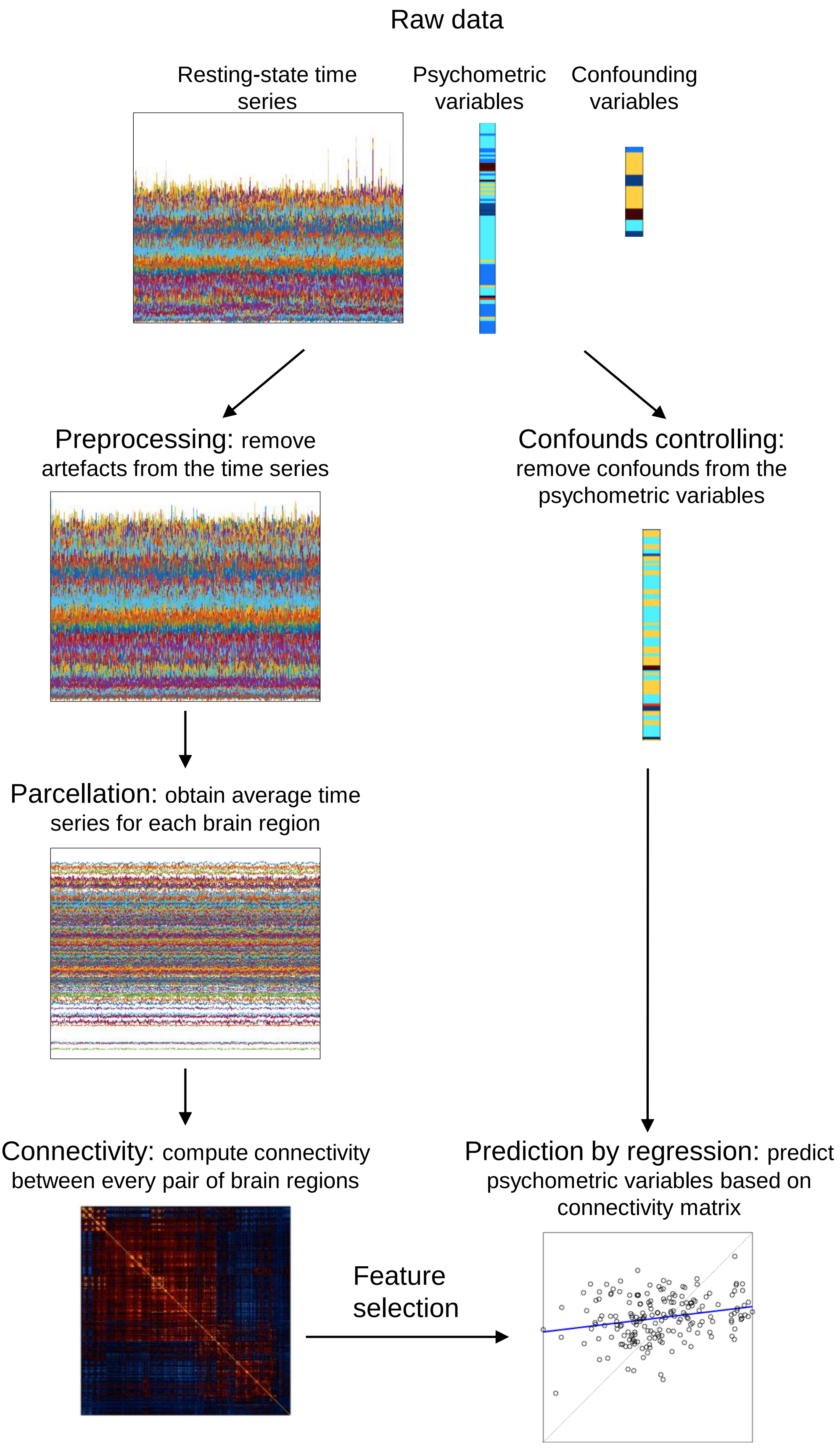


**Figure S1.** General workflow of a connectivity-based psychometric prediction (CBPP) framework

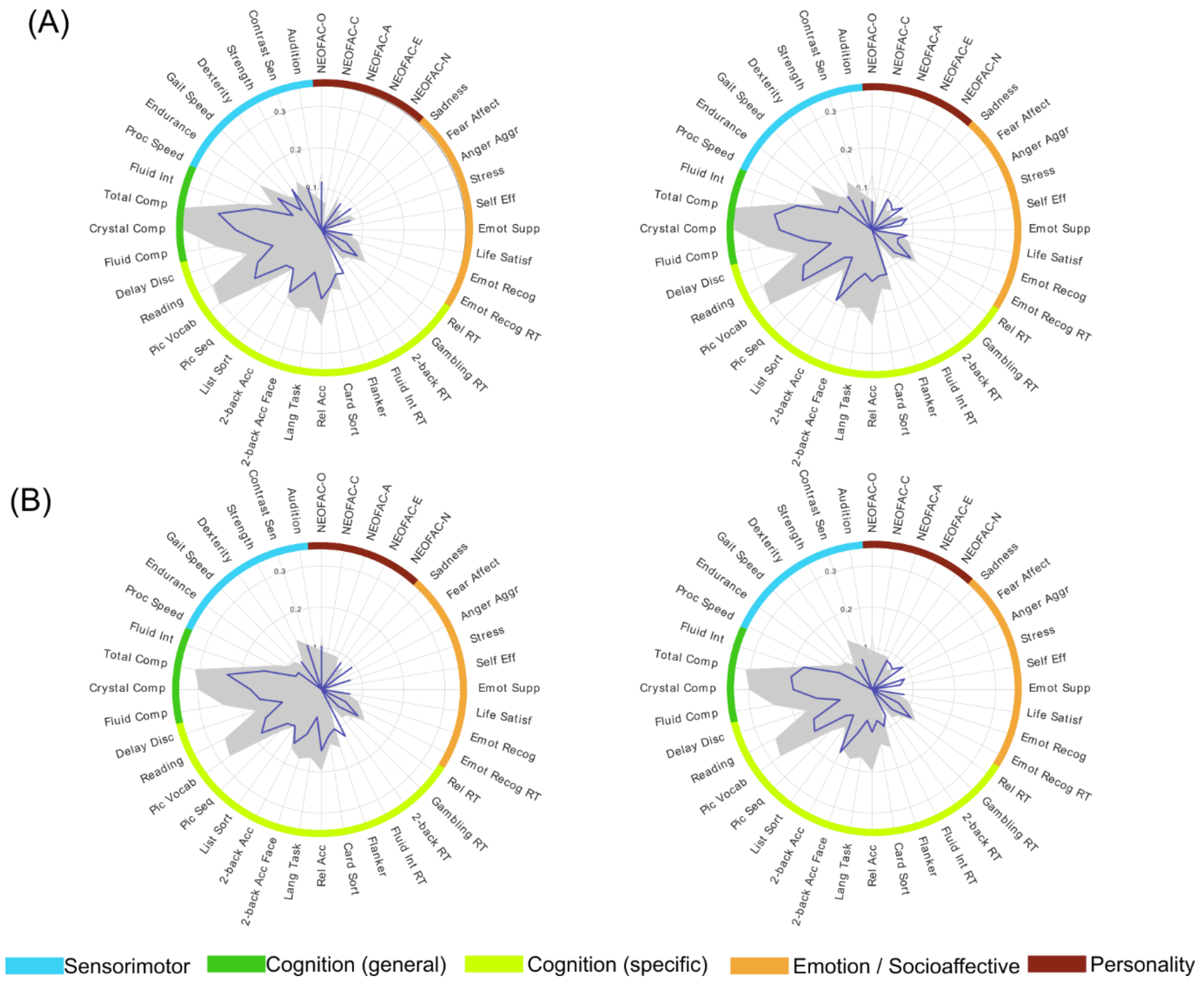


**Figure S2.** Psychometric profiles for pairs of parcels in supramarginal gyrus using (a) confounds excluding motion-related confounds and (b) confounds including motion-related confounds, in left and right hemispheres respectively. Gray filled contour shows whole-brain prediction profile, while blue contour shows parcel-wise prediction profile.

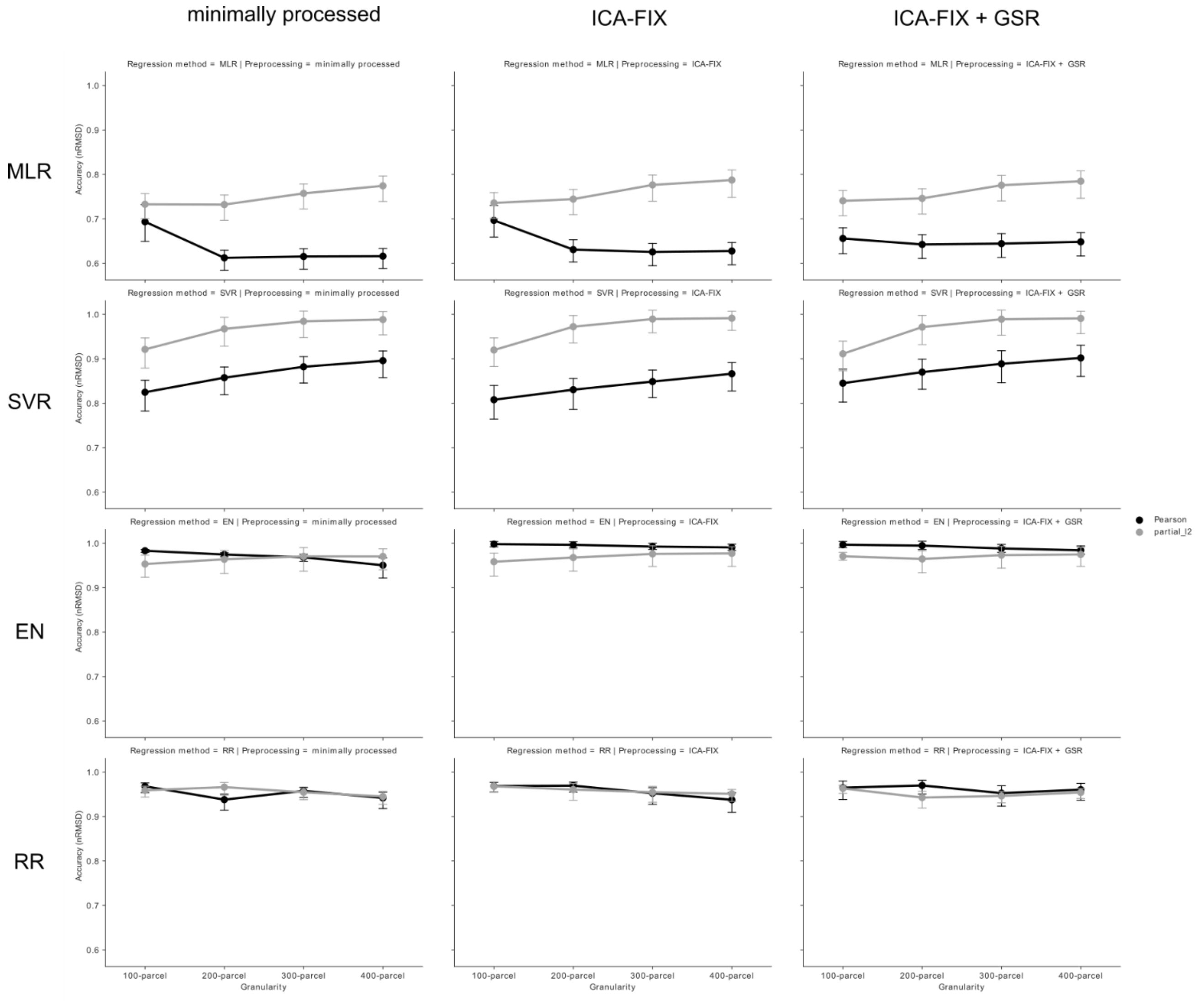


**Figure S3.** Average prediction accuracy (nRMSD between predicted and observed values) across the 40 psychometric variables for each combination of approaches in whole-brain CBPP. Error bars represent the 95% confidence interval across psychometric variables. The columns show combinations using minimally processed, FIX and FIX+GSR data respectively; the rows show combinations using MLR, SVR, EN and RR respectively. Dark lines represent combinations using Pearson correlation, while light grey lines represent combinations using partial correlation.

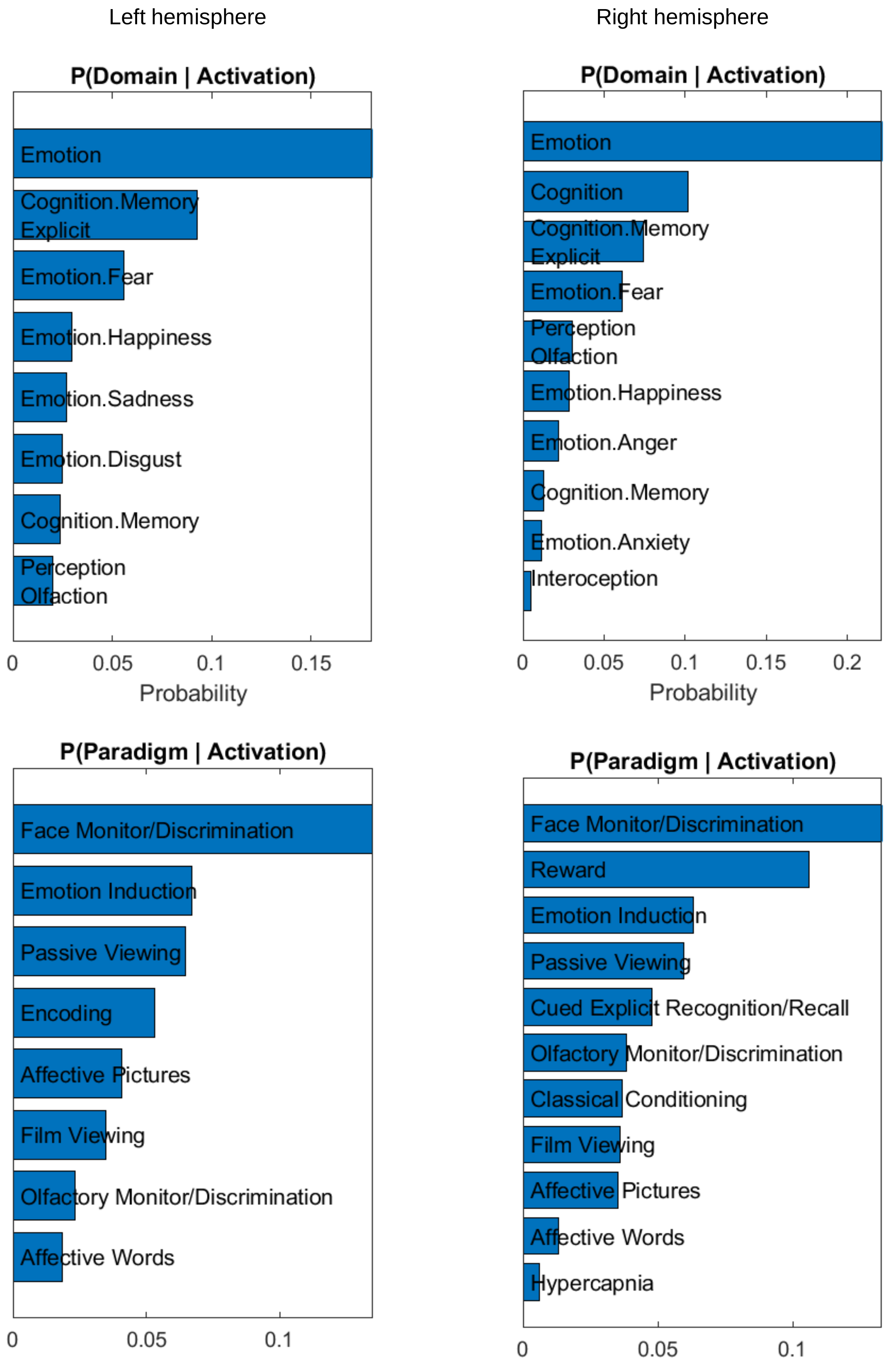


**Figure S4.** BrainMap behavioural profiles for the left and right anterior hippocampus parcels in AICHA atlas.

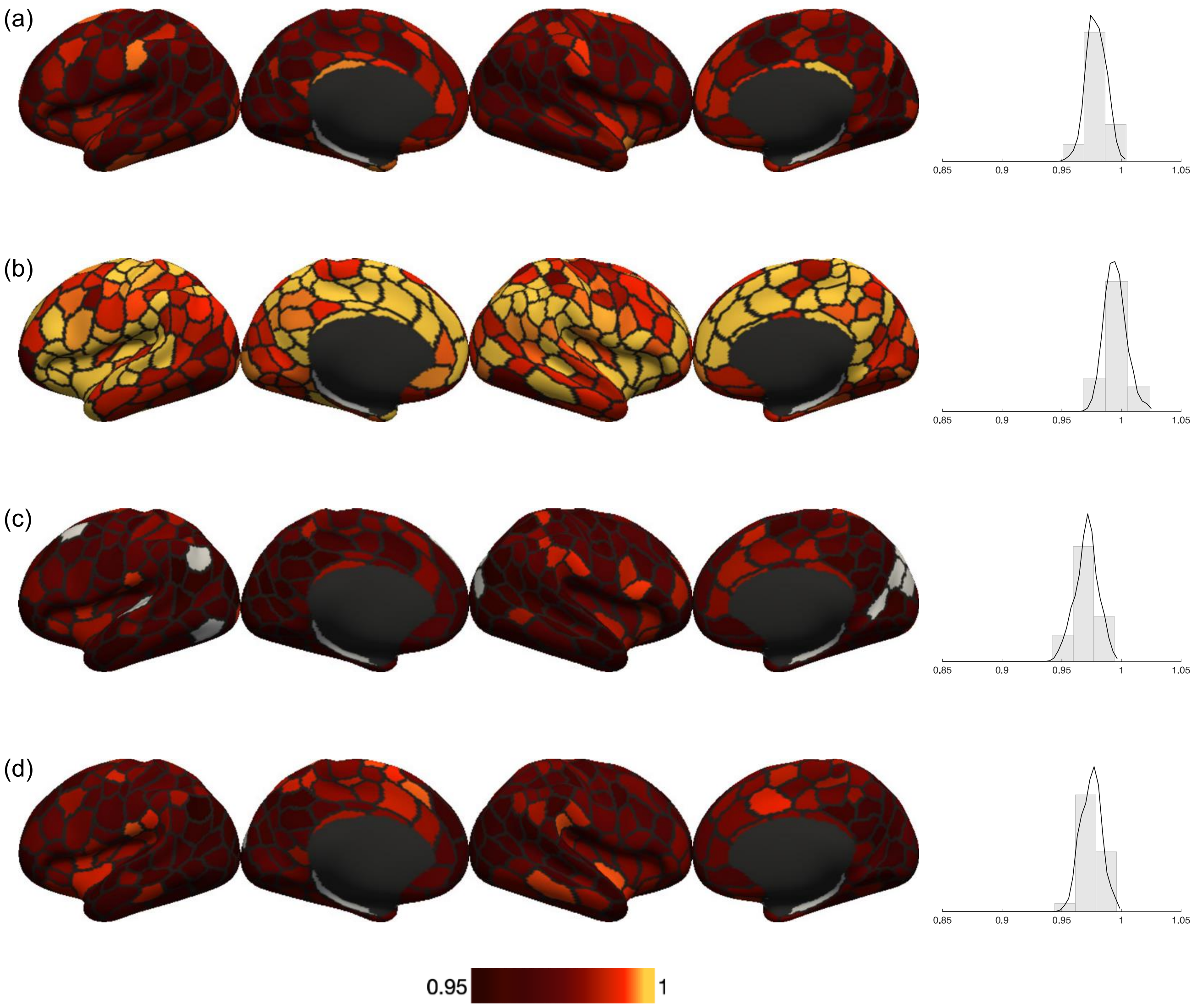


**Figure S5.** Prediction accuracy distribution using FIX-Pearson-SVR combination at 300-parcel granularity of the 4 selected psychometric variables: (a) strength (Strength) (b) crystallized cognition composite score (Crystal Comp) (c) working memory task overall accuracy (2-back Acc) (d) working memory task face condition accuracy (2-back Acc Face). Accuracies are normalized root mean squared deviation (nRMSD) between predicted and observed values. The rightmost column shows the histogram of prediction accuracies for each variable respectively.

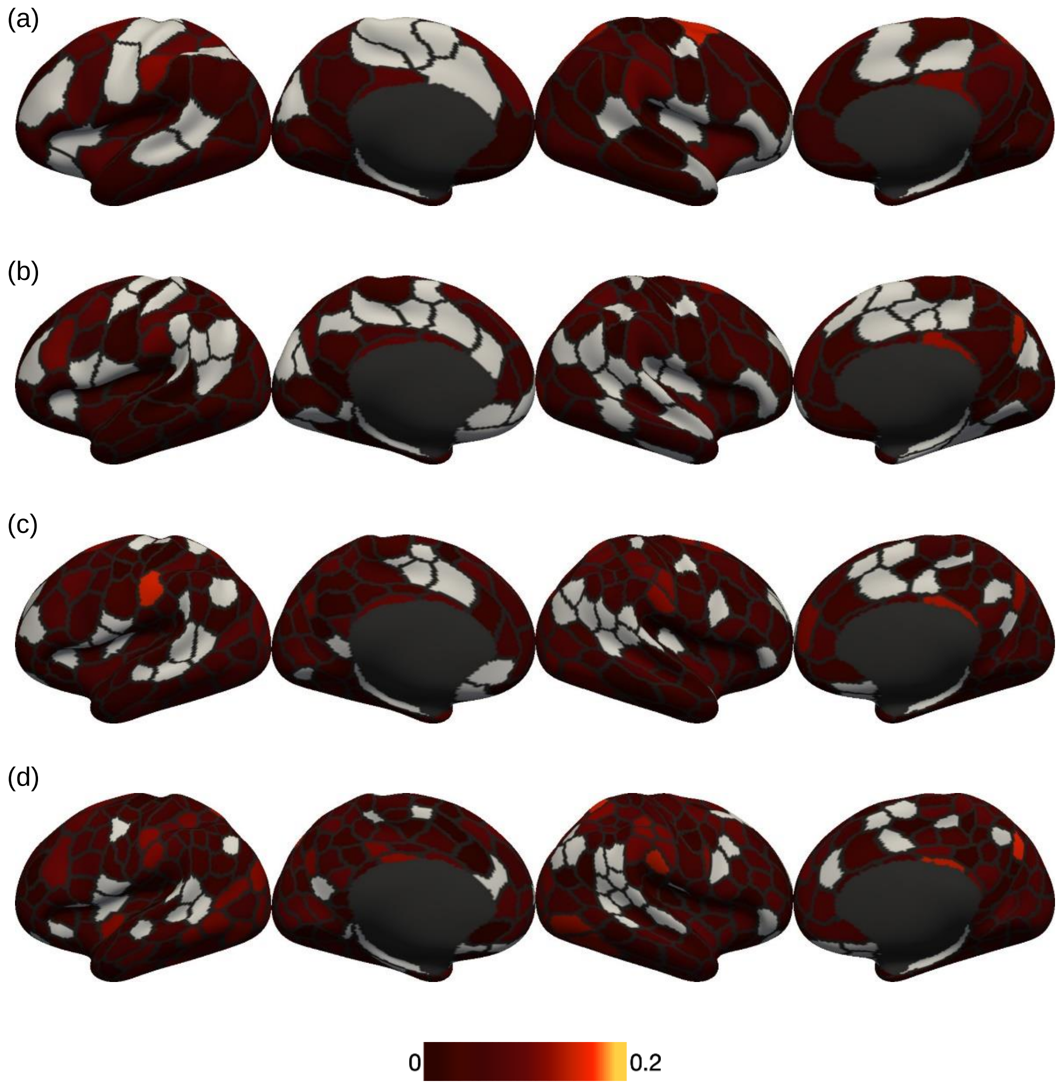


**Figure S6.** Prediction accuracy distribution using FIX-Pearson-SVR combination of strength (*Strength*) at (a) 100-parcel (b) 200-parcel (c) 300-parcel (d) 400—parcel granularity. Negative accuracies (i.e. correlation between predicted and actual values) were set to zeros and shown in gray.

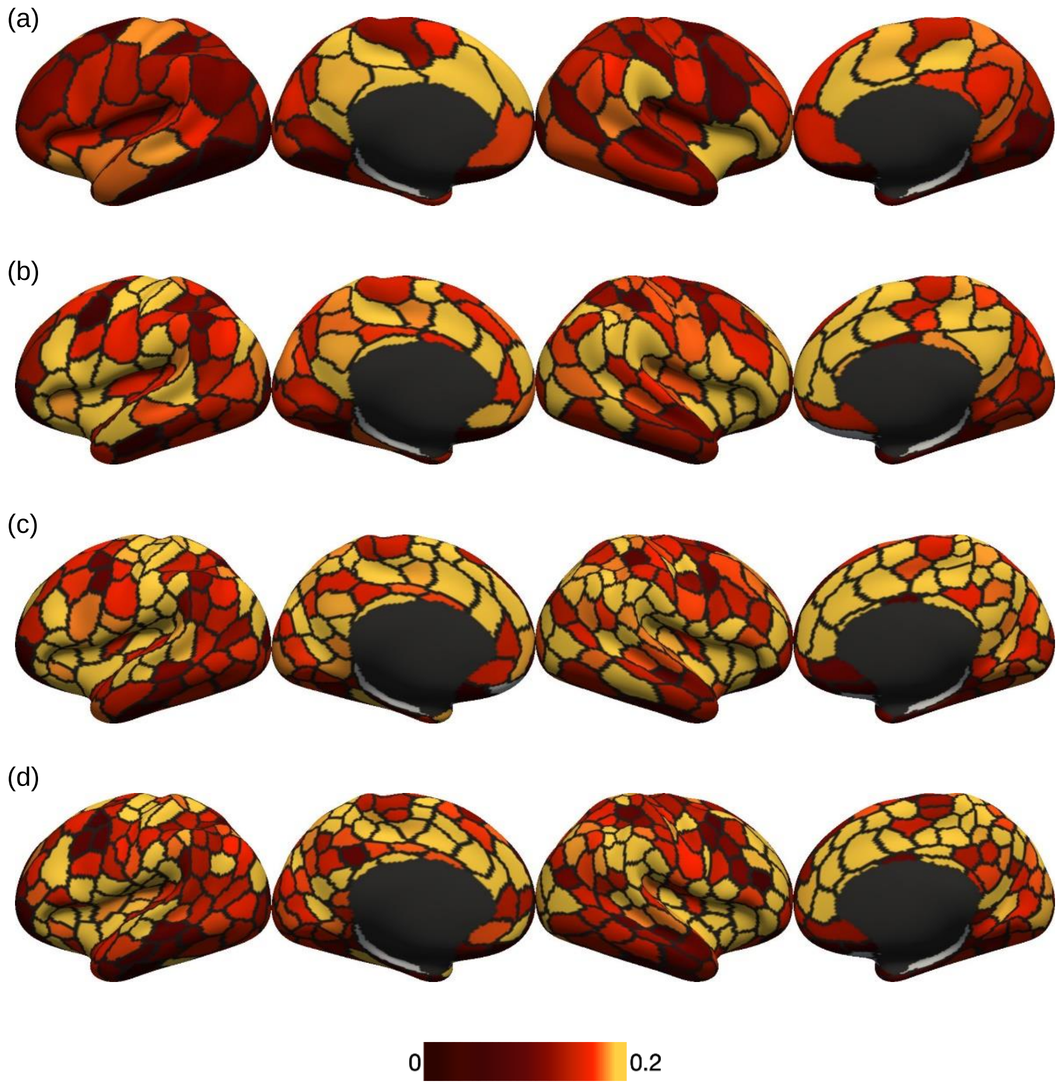


**Figure S7.** Prediction accuracy distribution using FIX-Pearson-SVR combination of crystallized cognition composite score (*Crystal Comp*) at (a) 100-parcel (b) 200-parcel (c) 300-parcel (d) 400—parcel granularity. Negative accuracies (i.e. correlation between predicted and actual values) were set to zeros and shown in gray.

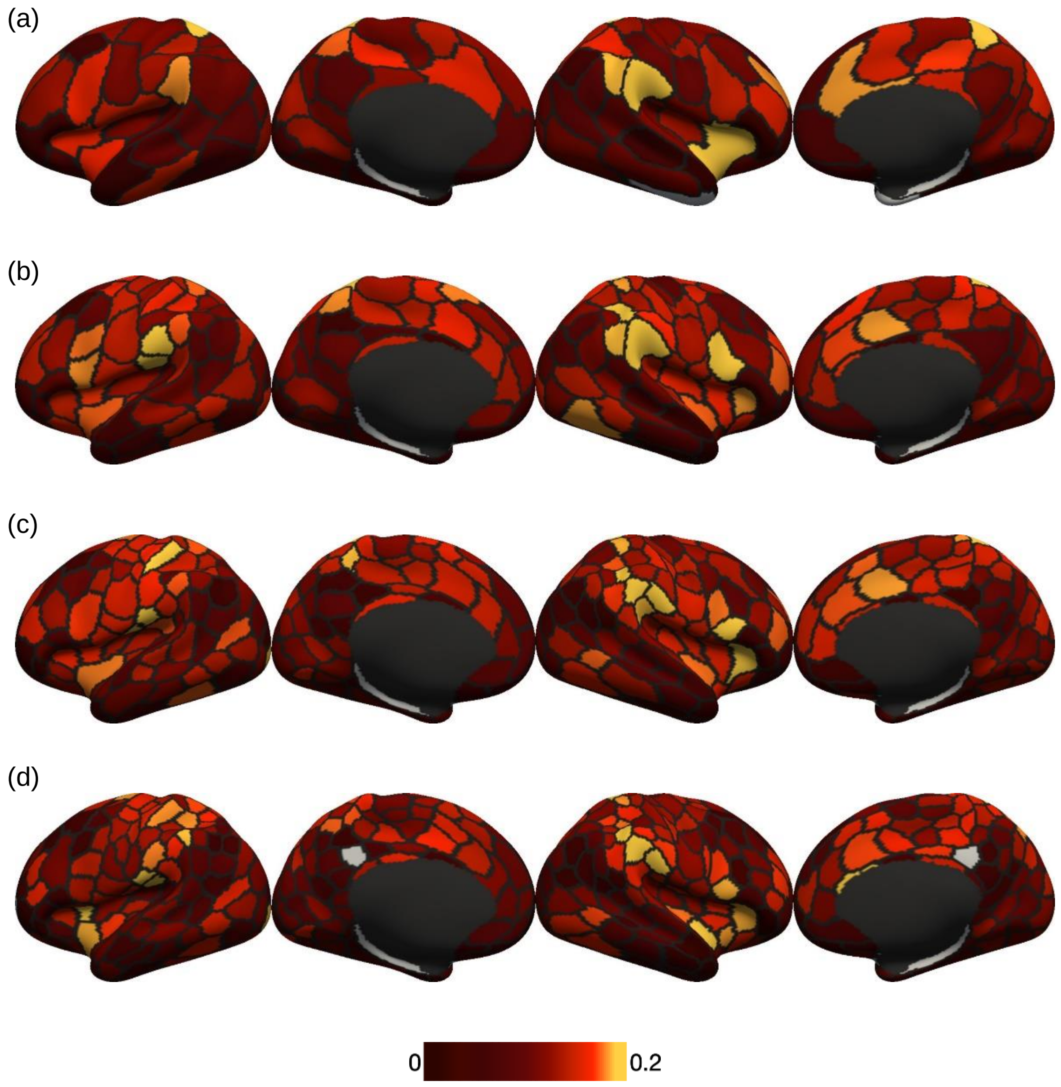


**Figure S8.** Prediction accuracy distribution using FIX-Pearson-SVR combination of working memory task overall accuracy (*2-back Acc*) at (a) 100-parcel (b) 200-parcel (c) 300-parcel (d) 400—parcel granularity. Negative accuracies (i.e. correlation between predicted and actual values) were set to zeros and shown in gray.

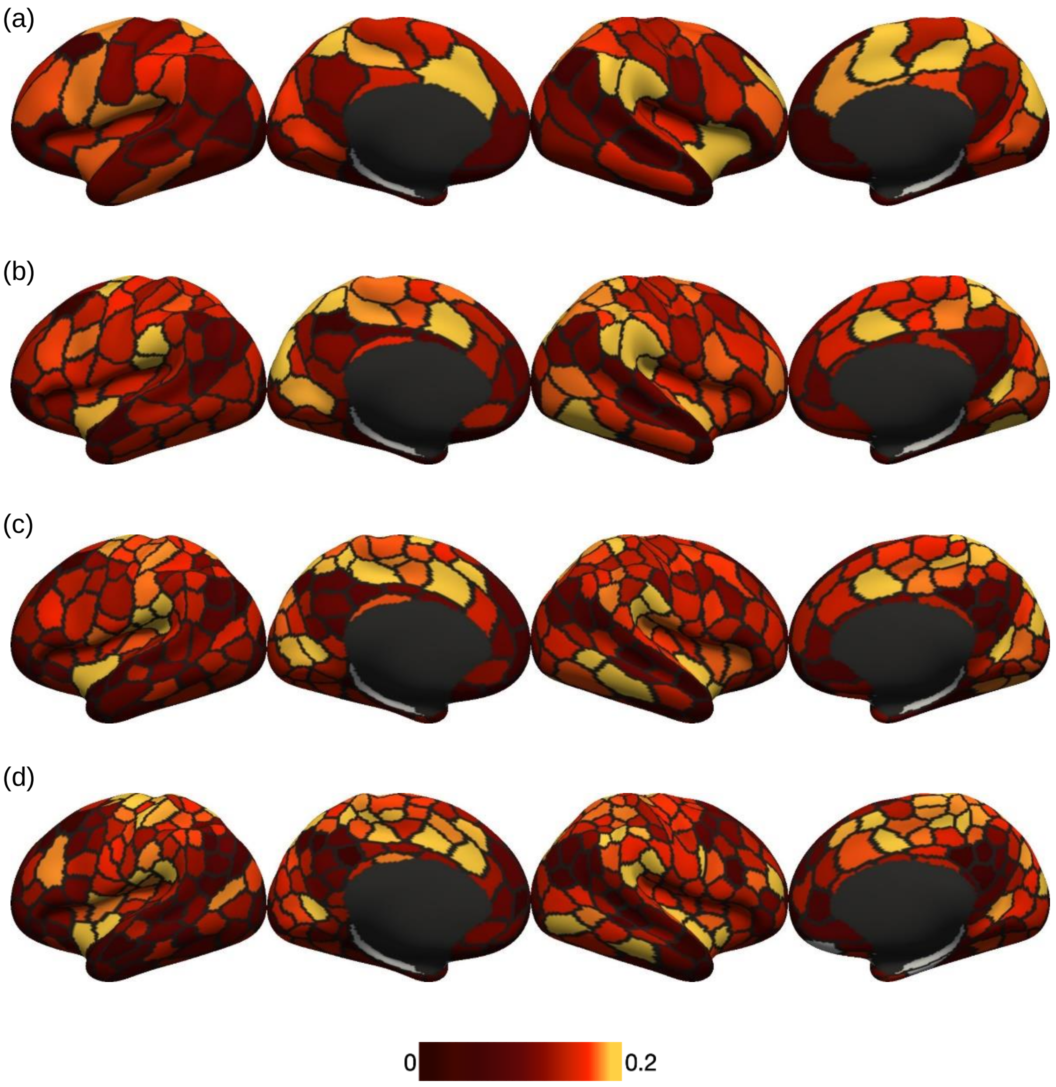


**Figure S9.** Prediction accuracy distribution using FIX-Pearson-SVR combination of working memory task face condition accuracy (*2-back Acc Face*) at (a) 100-parcel (b) 200-parcel (c) 300-parcel (d) 400—parcel granularity. Negative accuracies (i.e. correlation between predicted and actual values) were set to zeros and shown in gray.

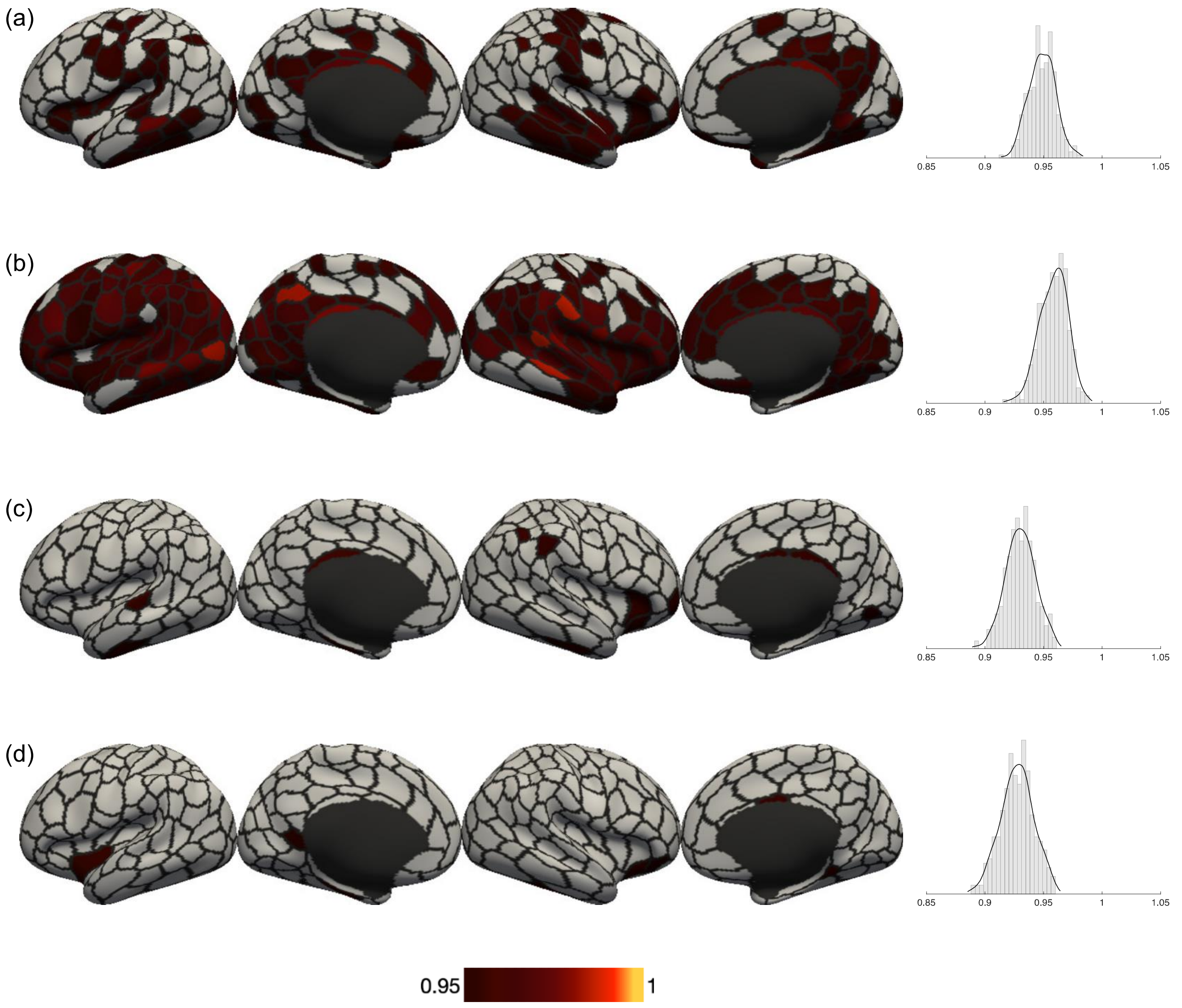


**Figure S10.** Prediction accuracy distribution using minimal-Pearson-SVR combination at 300-parcel granularity of the 4 selected psychometric variables: (a) strength (Strength) (b) crystallized cognition composite score (Crystal Comp) (c) working memory task overall accuracy (2-back Acc) (d) working memory task face condition accuracy (2-back Acc Face). Accuracies are normalized root mean squared deviation (nRMSD) between predicted and observed values. The rightmost column shows the histogram of prediction accuracies for each variable respectively.

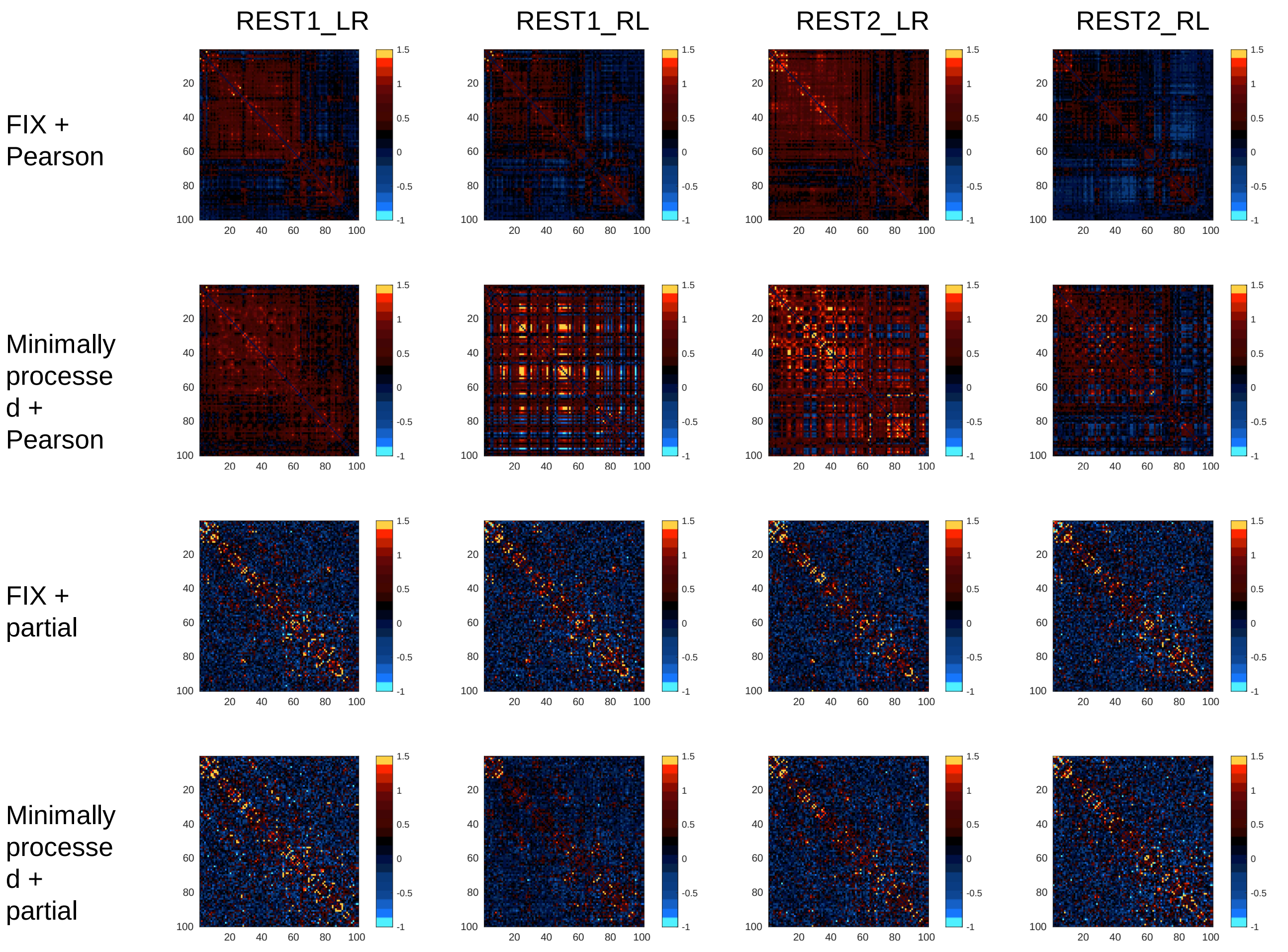


**Figure S11.** Functional connectivity matrix for the 4 runs of a single subject, using FIX data and minimally processed data, followed by Pearson correlation and partial correlation.

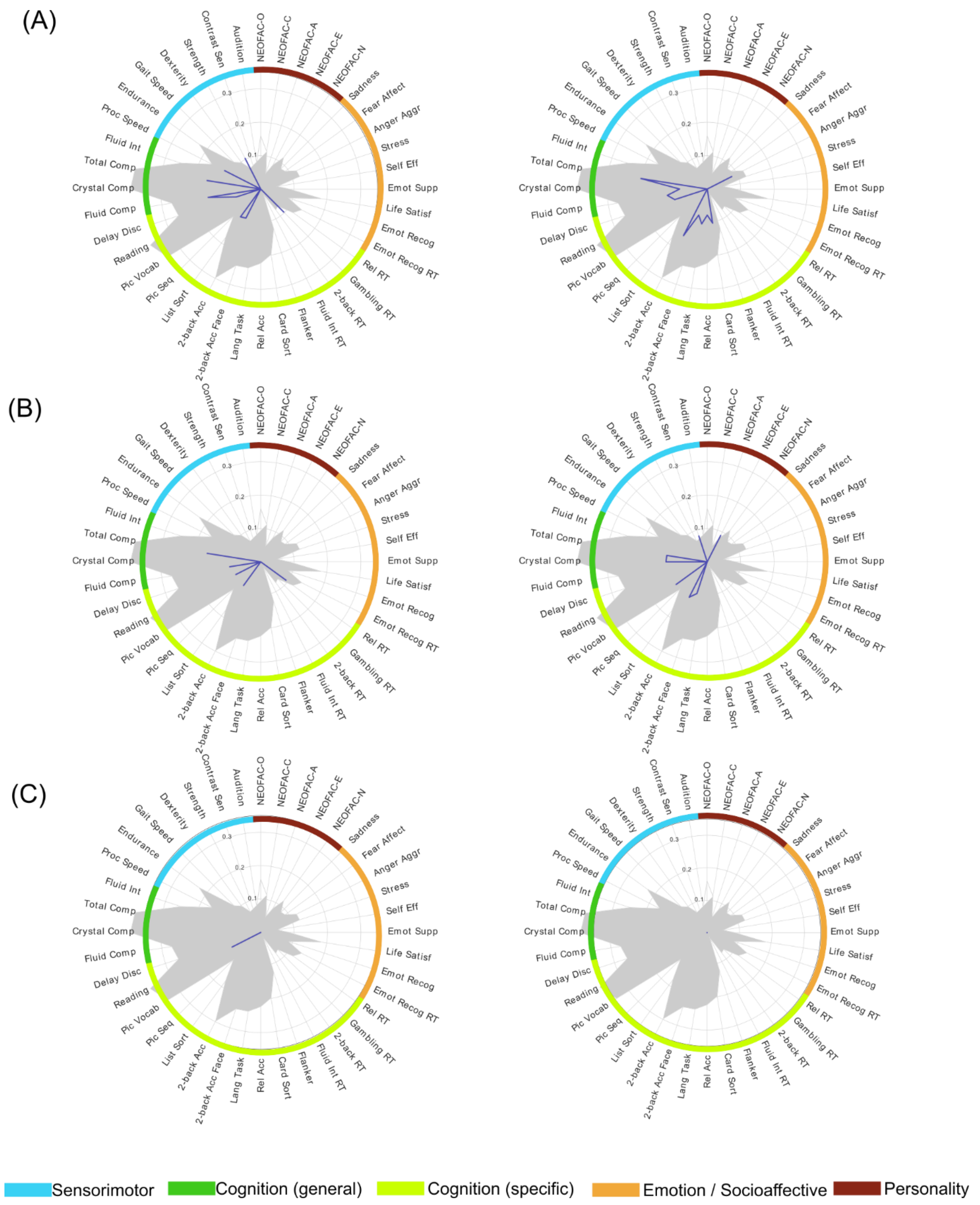


**Figure S12.** Psychometric profiles for pairs of parcels in (a) primary visual cortex (b) supramarginal gyrus and (c) Broca region in left and right hemispheres respectively, using FIX-partial-SVR combination at 300-parcel granularity, based on Pearson correlation accuracy. Gray filled contour shows whole-brain prediction profile, while blue contour shows parcel-wise prediction profile.

**Table S1.** The 40 selected psychometric variables

| Psychometric Variable | HCP Column Header |
|---|---|
| Audition | Noise_Comp |
| Contrast Sen | Mars_Final |
| Strength | Strength_AgeAdj |
| Dexterity | Dexterity_AgeAdj |
| Gait Speed | GaitSpeed_Comp |
| Endurance | Endurance_AgeAdj |
| Delay Disc | DDisc_AUC_40K |
| NEOFAC-O | NEOFAC_O |
| Crystal Comp | CogCrystalComp_AgeAdj |
| Reading | ReadEng_AgeAdj |
| Pic Vocab | PicVocab_AgeAdj |
| Pic Seq | PicSeq_AgeAdj |
| List Sort | ListSort_AgeAdj |
| 2-back Acc | WM_Task_2bk_Acc |
| 2-back Acc Face | WM_Task_2bk_Face_Acc |
| Lang task | Language_Task_Acc |
| Rel Acc | Relational_Task_Acc |
| Fluid Int | PMAT24_A_CR |
| Total Comp | CogTotalComp_AgeAdj |
| Fluid Comp | CogFluidComp_AgeAdj |
| Card Sort | CardSort_AgeAdj |
| Flanker | Flanker_AgeAdjk |
| Proc Speed | ProcSpeed_AgeAdj |
| Fluid Int RT | PMAT24_A_RTCR |
| 2-back RT | WM_Task_2bk_Median_RT |
| Gambling RT | Gambling_Task_Median_RT_Smaller |
| Rel RT | Relational_Task_Median_RT |
| Emot Recog RT | ER40_CRT |
| Emot Recog | ER40_CR |
| Life Satisf | LifeSatisf_Unadj |
| Emot Supp | EmotSupp_Unadj |
| Self Eff | SelfEff_Unadj |
| NEOFAC-E | NEOFAC_E |
| NEOFAC-A | NEOFAC_A |
| NEOFAC-C | NEOFAC_C |
| NEOFAC-N | NEOFAC_N |
| Stress | PercStress_Unadj |
| Anger Anggr | AngAggr_Unadj |
| Fear Affect | FearAffect_Unadj |
| Sadness | Sadness_Unadj |

**Table S2.** Pearson correlation between mean framewise displacement (FD) and psychometric variables

| Psychometric Variable (HCP column header) | Correlation with FD | Correlation with FD after regressing out the 9 confounds |
|---|---|---|
| Endurance_AgeAdj | 0.0941 | 0.0824 |
| DDisc_AUC_40K | 0.0803 | Not significant* |
| WM_Task_2bk_Face_Acc | 0.0782 | Not significant* |
| WM_Task_2bk_Acc | 0.0642 | Not significant* |
| ProcSpeed_AgeAdj | Not significant* | 0.0856 |
| CogFluidComp_AgeAdj | Not significant* | 0.0669 |

**Table S3.** Pearson correlation between DVARS and psychometric variables

| Psychometric Variable (HCP column header) | Correlation with DVARS | Correlation with DVARS after regressing out he 9 confounds |
|---|---|---|
| Strength_AgeAdj | 0.1187 | Not significant* |
| ProcSpeed_AgeAdj | 0.0898 | 0.0917 |
| CogTotalComp_AgeAdj | 0.0788 | 0.0799 |
| Language_Task_Acc | 0.0735 | 0.0709 |
| CogFluidComp_AgeAdj | 0.0687 | 0.0789 |
| Flanker_AgeAdj | 0.0655 | 0.0678 |
| FearAffect_Unadj | 0.0632 | Not significant* |
| Endurance_AgeAdj | Not significant* | 0.0712 |

* Significance for correlation values were assessed with Student's t distribution as null distribution (automatically by Matlab) with the threshold of $\alpha < 0.05$.

**Table S4.** Summary of existing psychometric prediction approaches in the CBPP framework

| Reference | Preprocessing | Parcellation granularity & Connectivity | Feature selection | Confounds | Regression |
|---|---|---|---|---|---|
| (Finn et al., 2015) | HCP minimal preprocessing pipeline, nuisance regression (motion, WM, CSV, and GS), detrending, LPF | 268x268 Pearson correlation matrix | Edges with significant correlation (P<0.01, P<0.05 and P<0.10) with target, summed according to sign (positive/negative) | Nil | Multiple linear regression with LOOCV |
| (Rosenberg et al., 2016) | MC, nuisance regression (motion, WM, CSF, GM), temporal smoothing | 268x268 Pearson correlation matrix | Edges with significant correlation (P<0.01) with target, summed according to sign | Nil | Multiple linear regression with LOOCV |
| (Smith et al., 2016) | HCP minimal preprocessing pipeline, ICA-FIX | Group-ICA of 15, 25, 50, 100, 200, 300 parcels; partial correlation with L2 regularization | Top 50% edges by correlation with targets | sex, age, $age^2$, sex*age, sex*$age^2$, brain & head size, overall head motion, acquisition quarter | Elastic net with leave-one-family-out cross-validation |
| (Noble et al., 2017) | HCP minimal preprocessing pipeline, nuisance regression (motion, WM, CSF, GS), detrending, LPF | 268x268 Pearson correlation matrix | Edges with significant correlation (P<0.05) with target, summed according to sign | Nil | Multiple linear regression with LOOCV |
| (Dubois et al., 2018) | HCP minimal preprocessing, followed by:<br>i) nuisance regression (motion, WM, CSF), LPF, GSR<br>ii) detrending, BPF, nuisance regression (motion, WM, CSF, GS) combined with censoring<br>iii) ICA-FIX, detrending, nuisance regression (GM, GS, CompCor of WM & CSF), censoring, BPF | i) 268x268 Pearson correlation matrix<br>ii) 360x360 Pearson correlation matrix | i) edges with significant correlation (P<0.01) with target, summed according to sign<br>ii) edges with significant correlation (P<0.01) with target | sex, age, handedness, fluid intelligence, brain size, motion, multiband reconstruction algorithm | Elastic net with leave-one-family-out cross-validation |
| (Li et al., 2019) | i) GSP data: STC, MC, nuisance regression (motion, WM, CSF, with/without GS), censoring, BPF<br>ii) HCP data: HCP minimal preprocessing, ICA-FIX, censoring, with/without GSR | 419x419 Pearson correlation matrix | Nil | sex, age, FD, DVARS | kernel ridge regression with 20-fold cross-validation |
| (Kashyap et al., 2019) | HCP minimal preprocessing pipeline, ICA-FIX, censoring, nuisance regression (motion, WM, CSF, GS), BPF | i) 419x419 Pearson correlation matrix<br>ii) 419x419 partial correlation matrix | Top 50% edges by correlation with targets | sex, age, FD | Elastic net with 20-fold cross-validation |

BPF, band-pass filtering; CSF, cerebrospinal fluid signal; DVARS, a summary score for data quality, defined as the spatial standard deviation of successive difference images; FD, frame-wise displacement; GM, gray matter signal; GS, global signal; GSR, global signal regression; GSP, Brain Genomics Superstruct Project; HCP: Human Connectome Project; LPF, low-pass filtering; MC, motion correction; STC, slice time correction; WM, white matter signal.

### Supplemental materials

#### 1. Whole-brain CBPP results (Pearson correlation accuracy)

For most combinations using partial correlation and some combinations using Pearson correlation with SVR or EN, prediction accuracies were generally observed to increase with granularity until 300-parcel level, before dropping at 400-parcel level ($p < 0.03$ corrected, $t > 2.202$, dof = 99). The trends were slightly different for MLR, where lower granularity usually show better performance than higher granularity ($p < 0.02$ corrected, $t > 2.365$, dof = 99). Almost no effect of granularity was found for combinations using RR.

The impact of connectivity computation methods was dependent on the preprocessing methods and regression methods used. For minimally processed data, all combinations using partial correlation significantly outperformed those using Pearson correlation ($p < 0.02$ corrected, $t > 2.365$, dof = 99). For other preprocessing methods, most combinations using SVR and all combinations using RR with FIX reached higher performance with partial correlation ($p < 0.01$ corrected, $t > 2.626$, dof = 99). On the other hand, all combinations using EN with FIX or FIX+GSR, at 400-parcel level, reached significantly higher performance with Pearson correlation ($p < 0.008$ corrected, $t > 2.707$, dof = 99).

Preprocessing methods comparisons showed that minimally processed data almost always led to worse results when Pearson correlation was used ($p < 0.02$ corrected, $t > 2.365$, dof = 99), but were mostly comparable to FIX and FIX+GSR data when partial correlation was used. The comparisons between FIX and FIX+GSR were mostly not statistically significant, except for combinations using RR with Pearson correlation, where FIX+GSR data always led to better results ($p < 0.001$ corrected, $t > 3.392$, dof = 99).

Finally, comparing the regression methods, SVR, EN and RR outperformed MLR in almost all combinations ($p < 0.001$ corrected, $t > 3.392$, dof = 99). Among these three regression methods, SVR and RR always outperformed EN when partial correlation was used ($p < 0.02$ corrected, $t > 2.365$, dof = 99), while SVR and EN outperformed RR when Pearson correlation and FIX data were used ($p < 0.001$ corrected, $t > 3.392$, dof = 99).

### Supplemental materials

#### 2. Whole-brain CBPP results (nRMSD accuracy)

Different trends of prediction accuracy by parcellation granularity were observed when different connectivity computation methods or regression methods were used. For combinations using partial correlation or SVR, prediction accuracies were generally observed to increase with granularity ($p < 0.02$ corrected, $t > 2.365$, dof = 99). Combinations using Pearson correlation generally show decrease in prediction accuracies with higher granularity for MLR and EN ($p < 0.03$ corrected, $t > 2.202$, dof = 99). No effect was observed in any combination using RR.

The impact of connectivity computation methods was dependent on the regression methods used. When MLR or SVR was used, combinations using partial correlation significantly outperformed those using Pearson correlation ($p < 0.001$ corrected, $t > 3.392$, dof = 99). On the other hand, for EN-FIX and EN-FIX+GSR combinations, higher performance was always achieved with Pearson correlation ($p < 0.01$ corrected, $t > 2.626$, dof = 99). Again, no effect was observed in any combination using RR.

Differing from correlation accuracy, preprocessing methods comparisons showed that minimally processed data only led to worse results when Pearson correlation and EN were used ($p < 0.003$ corrected, $t > 3.043$, dof = 99). The comparisons between FIX and FIX+GSR were mostly statistically significant for combinations using SVR, as FIX combinations showed higher performance when partial correlation was used ($p < 0.005$ corrected, $t > 2.871$, dof = 99), while FIX+GSR combinations showed higher performance when Pearson correlation was used ($p < 0.01$ corrected, $t > 2.626$, dof = 99). No effect was observed in combinations using RR.

Finally, comparing the regression methods, SVR, EN and RR outperformed MLR in all combinations ($p < 0.001$ corrected, $t > 3.392$, dof = 99). Among these three regression methods, EN and RR outperformed SVR in most combinations ($p < 0.03$ corrected, $t > 2.202$, dof = 99).
